# HIF-2α drives an intrinsic vulnerability to ferroptosis in clear cell renal cell carcinoma

**DOI:** 10.1101/388041

**Authors:** Yilong Zou, Michael J. Palte, Amy A. Deik, Haoxin Li, John K. Eaton, Wenyu Wang, Yuen-Yi Tseng, Rebecca Deasy, Maria Alimova, Vlado Dančík, Elizaveta S. Leshchiner, Vasanthi S. Viswanathan, Sabina Signoretti, Toni K. Choueiri, Jesse S. Boehm, Bridget K. Wagner, John Doench, Clary B. Clish, Paul A. Clemons, Stuart L. Schreiber

## Abstract

Kidney cancers are characterized by extensive metabolic reprogramming and resistance to a broad range of anti-cancer therapies. By interrogating the Cancer Therapeutics Response Portal compound sensitivity dataset, we show that cells of clear-cell renal cell carcinoma (ccRCC) possess a lineage-specific vulnerability to ferroptosis that can be exploited by inhibiting glutathione peroxidase 4 (GPX4). Using genome-wide CRISPR screening and lipidomic profiling, we reveal that this vulnerability is driven by the HIF-2α–HILPDA pathway by inducing a polyunsaturated fatty acyl (PUFA)-lipid-enriched cell state that is dependent on GPX4 for survival and susceptible to ferroptosis. This cell state is developmentally primed by the HNF-1β–1-Acylglycerol-3-Phosphate O-Acyltransferase 3 (AGPAT3) axis in the renal lineage. In addition to PUFA metabolism, ferroptosis is facilitated by a phospholipid flippase TMEM30A involved in membrane topology. Our study uncovers an oncogenesis-associated vulnerability, delineates the underlying mechanisms and suggests targeting GPX4 to induce ferroptosis as a therapeutic opportunity in ccRCC.

**HIGHLIGHTS:**

- ccRCC cells exhibit strong susceptibility to GPX4 inhibition-induced ferroptosis
- The GPX4-dependent and ferroptosis-susceptible state in ccRCC is associated with PUFA-lipid abundance
- The HIF-2α–HILPDA axis promotes the selective deposition of PUFA-lipids and ferroptosis susceptibility
- AGPAT3 selectively synthesizes PUFA-phospholipids and primes renal cells for ferroptosis

## INTRODUCTION

Despite significant surgical and systemic therapy advances in kidney cancers, the recurrent and metastatic form of kidney malignancies remains largely fatal, with incidences increasing worldwide [1, 2]. Although cancers arising from different segments of the renal tubule display distinct histological features and clinical courses, genetic mutations present in various kidney cancers are highly clustered in metabolic pathways that regulate oxygen, iron, nutrient, and energy homeostasis [3–5]. The extensive metabolic reprogramming underscores a common metabolic state that kidney cancer cells acquire during tumorigenesis. One emerging theme of this cell state is dysregulated lipid metabolism. Clear cell renal cell carcinoma (ccRCC), the most frequent form of RCC, features by aberrant accumulation of glycogen and lipid droplets [6–9], whereas renal oncocytomas exhibit early loss of mitochondrial complex I genes, and stalled lipid β-oxidation [10, 11]. While these metabolic alterations support cancer progression and promote resistance to immune surveillance and therapies [5, 12], directly targeting the primary metabolic liabilities remains challenging due to complications from metabolic plasticity and systematic toxicity of available compounds [13, 14]. Exploring novel vulnerabilities associated with the metabolic states found in kidney cancer cells is critical for developing new therapies.

Recently, we generated quantitative sensitivity profiles of 481 “Informer Set” compounds in 887 cancer cell lines from various lineages, including RCC, and made these data and analysis tools available in the Cancer Therapeutics Response Portal (CTRP) [15–17]. These compounds perturb many distinct nodes in cellular pathways, and therefore are informative for identifying both pancancer and tissue-specific vulnerabilities. Here, by interrogating the CTRP, we report that inhibitors of glutathione peroxidase 4 (GPX4), among all nodes of cell circuitry, exhibit the highest selectivity and potency in killing RCC cells, which are largely represented by the ccRCC subtype. ccRCC is intrinsically refractory to chemotherapy and radiation therapy and highly prone to metastasis, presenting an unmet medical need [18–20]. ccRCC cells gain constitutive activation of hypoxia-inducible factor-2a (HIF-2α) through inactivation of *von Hippel-Lindau (VHL)* tumor suppressor [21, 22]. HIF-2α promotes glycolysis and angiogenesis, suppresses oxidative phosphorylation, and drives lipid accumulation [8, 23, 24]. The broad metabolic alterations driven by HIF-2α have been translated into diagnosis tools and biomarkers for ccRCC. However, therapeutic strategies targeting HIF-2α and its downstream effectors have been less successful, with modest responses and emergence of resistance in experimental cancer models [6, 25, 26], though recently a first-in-class HIF-2α antagonist showed activity and a favorable toxicity profile in a heavily pretreated RCC population [27]. While the genetic etiology of ccRCC has been illuminated, the lack of actionable targets underscores the pressing need for novel biological insights. Here, our CTRP analyses, along with previous characterizations, link ccRCC to a unique dependency on GPX4 [28, 29]. Given GPX4’s role in selectively detoxifying lipid hydroperoxides, and that GPX4 inhibition triggers ferroptotic cell death (ferroptosis) [28], we postulate that ccRCC cells are intrinsically susceptible to ferroptosis, and targeting GPX4 represents a therapeutic opportunity in this devastating disease.

Though the developmental role of ferroptosis has not yet been identified, ferroptotic death is associated with various pathological conditions, including acute kidney injury, hepatocellular degeneration and hemochromatosis, traumatic brain injury and neurodegeneration [30–36]. Notably, we and others have recently identified a GPX4-dependent state in cancer cells that in general resists apoptotic death otherwise induced by the main modalities of cancer treatments – chemotherapy, targeted therapy, and immunotherapy [37–39]. These insights point to ferroptosis-modulatory agents as attractive therapeutic strategies for cancer treatment. However, the susceptibility to ferroptosis varies significantly among organ systems and the mechanisms underlying cell type-specific ferroptosis sensitivity are poorly understood.

In the present study, we systematically characterize the mechanisms driving the lineage-specific GPX4 dependency and ferroptosis susceptibility in ccRCC. By combining genome-wide CRISPR screening and lipidomic profiling, we identify HIF-2α as a central driver of this vulnerability. HIF-2α selectively enriches polyunsaturated fatty acyl (PUFA)-lipids, and creates a ferroptosis-susceptible cell state. Our findings have implications for understanding the mechanisms of the ferroptosis pathway, as well for developing novel treatment options for ccRCC patients.

## RESULTS

### ccRCC cells are intrinsically susceptible to GPX4 inhibition-induced ferroptosis

In search of druggable vulnerabilities for kidney cancers, we interrogated the CTRP (<u>portals.broadinstitute.org/ctrp/</u>) chemical sensitivity dataset to identify compounds that selectively inhibit the survival and proliferation of kidney cancer cells [16, 17]. The kidney-derived cancer cell lines in CTRP included 17 ccRCC, 1 renal transitional-cell carcinoma (RTCC) and 2 other cell lines with uncertain subtype classification [40, 41] (**Figure S1A**). Among the 481 CTRP compounds that selectively target a broad array of biological pathways, commonly used chemotherapeutic agents, such as paclitaxel, exhibited the lowest potency in ccRCC cells compared with other solid cancer cell lines (sCCL), indicating a tendency towards resistance to apoptosis-inducing agents in ccRCC (**Figure 1A**). In contrast, three GPX4 inhibitors emerged as the most potent and selective compounds for killing ccRCC cells: (1S,3R)-RSL3 (RSL3), ML210 and, ML162 (**Figures 1A-B and S1A**) [28, 38, 42]. Selective killing by inhibitors of GPX4 strongly indicates that ccRCC cells are vulnerable to ferroptosis. Two other compounds, GSK-3β inhibitor ML320 and PI3K inhibitor GSK-2636771, also exhibited selectivity for ccRCC cells but with moderate sensitivity (**Figure 1A**) [43, 44]. Grouping cancer cells by tissue-of-origin revealed that kidney cancer cells are most sensitive to GPX4 inhibitors among solid cancers, while being the most insensitive to paclitaxel (**Figures 1C-E and S1B**). Collectively, our systematic analyses phenocopy the intrinsic resistance to chemotherapies and modest response to targeted therapies in ccRCC patients, and suggest that GPX4 is a targetable dependency in this difficult-to-treat disease.

**Figure 1.**
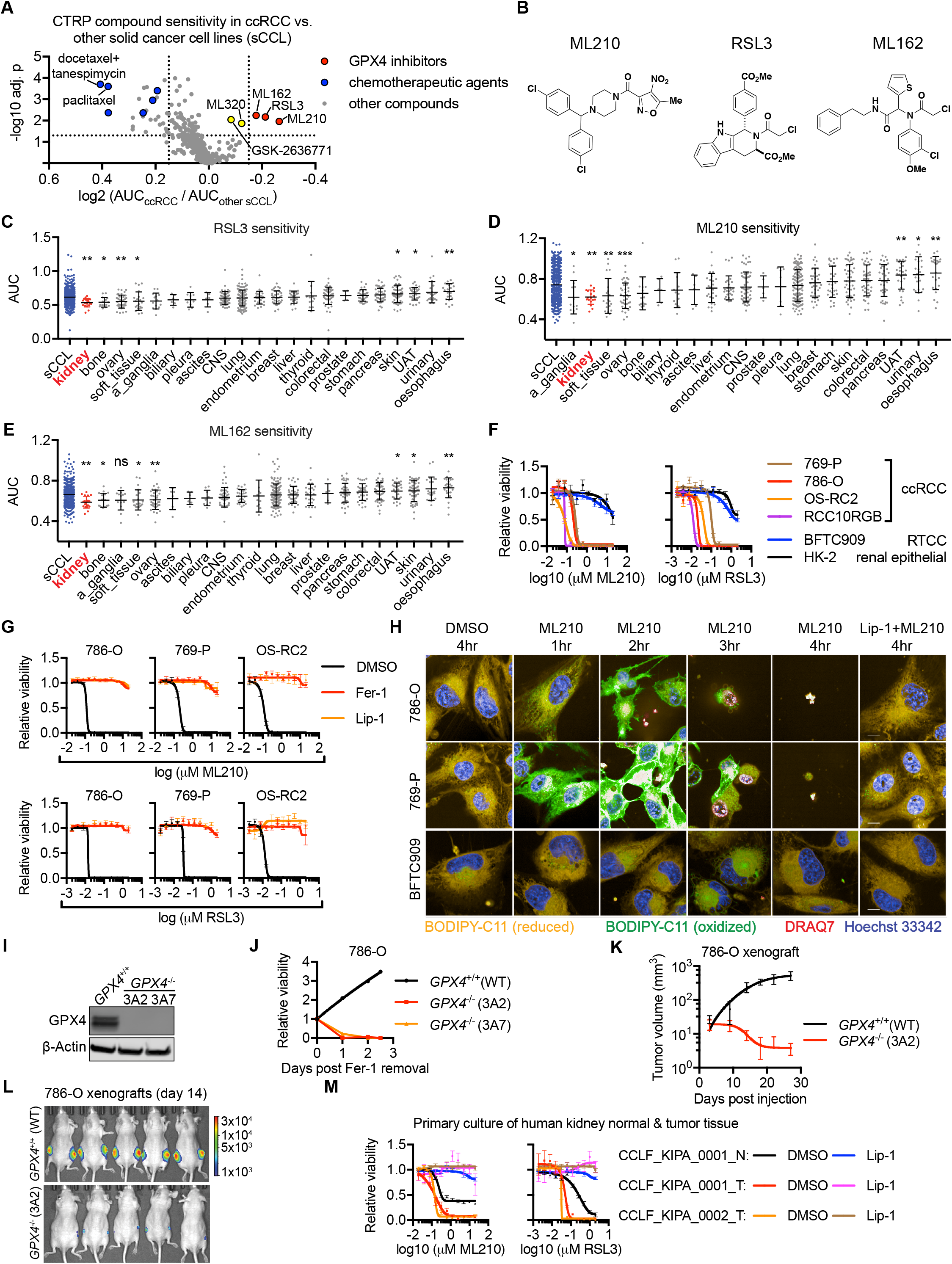
ccRCC cells are intrinsically vulnerable to GPX4 inhibition-induced ferroptosis. **A.** Volcano plot showing compound sensitivity comparison by normalized area-under-curve (AUC) values between clear cell renal cell carcinoma (ccRCC) cells (N=17) and other solid tumor cancer cell lines (sCCL) (N=643) in the Cancer Therapeutics Response Portal (CTRP). Log-ratios of the average AUC for ccRCC versus other sCCL cells are plotted against the significance of a Mann-Whitney-Wilcoxon test performed between the two distributions for each compound (represented by each dot). Compounds to the left exhibit lower sensitivity in ccRCC cells than other sCCL, whereas compounds to the right exhibit higher sensitivity in ccRCC cells. ML162, ML210, RSL3: GPX4 inhibitors. ML320, a GSK-3β inhibitor. GSK-2636771, a PI3K inhibitor. Blue dots: chemotherapeutic agents. **B.** Chemical structures of GPX4 inhibitors ML210, RSL3 and ML162. **C.** Scatterplot of AUC value distributions for RSL3 in sCCL (blue) or cancer cell lines from each specified tissue-of-origin, including kidney (orange). Larger AUC values indicate lower compound sensitivity and vice versa. Tissue types are ordered by the average AUC values. Abbreviations for tissue-of-origin: CNS, central nervous system; UAT, upper aerodigestive tract; a_ganglia, autonomic ganglia. A Mann-Whitney-Wilcoxon test was performed between each tissue and sCCLs from other tissues. Statistical significance is adjusted for multi-test correction. *, p<0.05; **, p<0.01. Line and error bars in each group represent mean and standard deviation (S.D.). **D.** Scatterplot of AUC value distributions for GPX4 inhibitor ML210 in sCCL or cell lines from each specified tissue-of-origin. ***, p<0.001; ns, not significant. **E.** Scatterplot of AUC value distributions for GPX4 inhibitor ML162 in sCCL or cell lines from each specified tissue-of-origin. **F.** Viability curves for ccRCC cell lines 769-P, 786-O, OS-RC2, and RCC10RGB, renal transitional cell carcinoma (RTCC) cell line BFTC909, and immortalized, non-transformed normal renal proximal tubule epithelial cell lines HK-2 under a 11-point, 2-fold dilution series of ML210 (left) or RSL3 (right) treatment. **G.** Viability curves for 786-O, 769-P, and OS-RC2 cells treated with ML210 or RSL3 plus indicated DMSO, liproxstatin-1 (Lip-1) or ferrostatin-1 (Fer-1). **H.** Confocal images of 786-O, 769-P, and BFTC909 cells treated with ML210 plus DMSO or Lip-1 for the indicated time periods. Oxidation of the BODIPY-C11 dye (turning from orange to green) was employed to report lipid reactive oxygen species (ROS) levels. DRAQ7 and Hoechst 33342 reports dying or live cells, respectively. Scale bars in the images represent 10μm. **I.** Immunoblot analysis of GPX4 protein levels in wildtype (GPX4^+/+^) 786-O or two *GPX4^−/−^* single-cell clones #3A2 and #3A7. β-Actin was used as a loading control. **J.** Viability curves for wildtype *(GPX4^+/+^)* 786-O or two *GPX4^−/−^* single-cell clones #3A2 and #3A7 throughout a 2.5-day time course after Fer-1 removal. Viability is normalized to that at day 0 of Fer-1 removal. **K.** Tumor volume measurements of xenografts derived from subcutaneous injection of wildtype (GPX4^+/+^) 786-O or *GPX4^−/−^* 3A2 cells (five mice per group, two tumors per mouse). Matrigel containing 0.5μM Lip-1 was used to assist the initial implantation of the cancer cells. **L.** Representative images of luciferase imaging in tumors at day 14 post implantation as shown in panel K. **M.** Viability curves for primary human ccRCC cell lines CCLF_KIPA_0001_T, CCLF_KIPA_0002_T, and normal renal-cell culture CCLF_KIPA_0001_N from the same patient as CCLF_KIPA_0001_T treated with a 11-point, 2-fold dilution series of ML210 (left) or RSL3 (right), plus DMSO or 0.5μM of Lip-1. For viability assays in panel **F,G,J** and **M**, viability under each condition is relative to that of the respective DMSO-treated condition. Each data point has four biological replicates, and error bars represent ±S.D. Representative plot of experiments repeated three times is presented. See also Figure S1.

To validate the GPX4 dependency in ccRCC, we assessed the sensitivity to GPX4 inhibitors in four frequently used ccRCC cell lines in CTRP: 786-O, 769-P, OS-RC2, and RCC10RGB. Compared with two control cell lines, immortalized renal tubule cell line HK-2 and RTCC line BFTC909, all four ccRCC cell lines were more sensitive to GPX4 inhibitors (**Figure 1F**). These results are consistent with previous characterization of the sensitivity to glutathione depletion or treatment with erastin, a system Xc^-^ *(SLC7A11)* inhibitor and a moderate ferroptosis inducer, within the NCI-60 cancer cell lines [28, 29]. shRNA-mediated *GPX4* knockdown significantly reduced the viability of 786-O, 769-P and OS-RC2 cells (**Figure S1C**), suggesting a genetic dependency on *GPX4.* We verified these results by interrogating the complementary (to CTRP) Cancer Dependency Map (DepMap, https://depmap.org/portal) database for systematic characterizations of gene essentiality via genetic perturbations [45, 46]. In DepMap, both CRISPR and RNAi-based perturbations confirmed a strong dependency on *GPX4* in kidney cancer cells (**Figures S1D-E**). To determine whether ferroptosis is induced by GPX4 inhibition in ccRCC cells, we tested the activities of ferroptosis rescue agents ferrostatin-1 (Fer-1) and liproxstatin-1 (Lip-1), which act as lipophilic anti-oxidants to protect cells from oxy- and peroxy-radicals that derive from lipid hydroperoxides [47–49]. Indeed, Fer-1/Lip-1 treatment rescued the cell-killing effects of GPX4 inhibitors and blocked the accumulation of lipid reactive oxygen species (ROS), as determined by oxidized BODIPY-C11 levels (**Figures 1G-H and S1F**), confirming the induction of ferroptosis in ccRCC cells by GPX4 inhibitors.

We next generated xenografts derived from GPX4-depleted cells to evaluate the GPX4 dependency of ccRCC tumors *in vivo.* Single-cell 786-O clones with biallelic *GPX4* knockout were generated by CRISPR/Cas9 in the presence of Fer-1, where Fer-1 withdrawal triggers rapid cell death, confirming the essentiality of GPX4 in 786-O cells *in vitro* (**Figures 1I-J**). *In vivo, GPX4^−/−^* tumors regressed rapidly after implantation, in contrast to the continuous growth of *GPX4^+/+^* tumors (**Figures 1K-L**). This result confirms that ccRCC tumors are dependent on GPX4 *in vivo.* We then tested the sensitivity to GPX4 inhibitors in patient-derived primary cancer cells. CCLF_KIPA_0001_T and CCLF_KIPA_0002_T, two VHL-deficient, HIF-2α-positive primary ccRCC cell lines, exhibited significantly higher sensitivity to GPX4 inhibition than CCLF_KIPA_0001_N, a normal kidney tissue culture from the same patient as CCLF_KIPA_0001_T (**Figures 1M and S1G**). Lip-1 treatment blocked GPX4 inhibitor-induced cell death in all primary cells, confirming the involvement of ferroptosis (**Figure 1M**). Our data indicate that human ccRCC tumors are highly susceptible to GPX4 inhibition-induced ferroptosis, and a sensitivity window is potentially present between tumor and normal renal tissues.

### Genome-wide CRISPR resistance screen identifies novel mediators for ferroptosis susceptibility in ccRCC cells

*GPX4* is ubiquitously expressed at largely constant levels across all major cancer types in The Cancer Genome Atlas (TCGA) and CTRP databases [50, 51], and thus GPX4 dependency appears independent of *GPX4* expression levels (**Figures S2A-B**). To explore mechanisms driving the lineage-specific GPX4 dependency in ccRCC, we performed a genome-wide CRISPR resistance screen to identify mediators of ML210 sensitivity (**Figure 2A**). 786-O-Cas9 cells were transduced with a genome-wide library of 77,441 sgRNAs [52], treated with DMSO or ML210, and sequenced for barcode/sgRNA identification (**Figures 2A and S2C**). Collectively, 15 target genes were significantly enriched in all ML210-treated conditions (**Figure S2D**). The top hits included acyl-CoA synthetase long-chain family member 4 *(ACSL4)* and Kelch-like ECH associated protein 1 *(KEAP1)* (**Figures 2B and S2E**), two genes previously implicated as requirements for ferroptosis [38, 53–57]. Doxycycline-inducible sgRNA expression in Cas9-expressing 786-O and 769-P cells further verified ACSL4 and KEAP1 as necessary for compound-induced ferroptosis (**Figures 2C-F and S2C**). In addition, KEAP1-depletion induced accumulation of NRF2 *(NFE2L2)* protein (**Figure 2E**), suggesting the mitigation of ferroptosis by activation of anti-oxidant pathways [54, 57].

**Figure 2.**
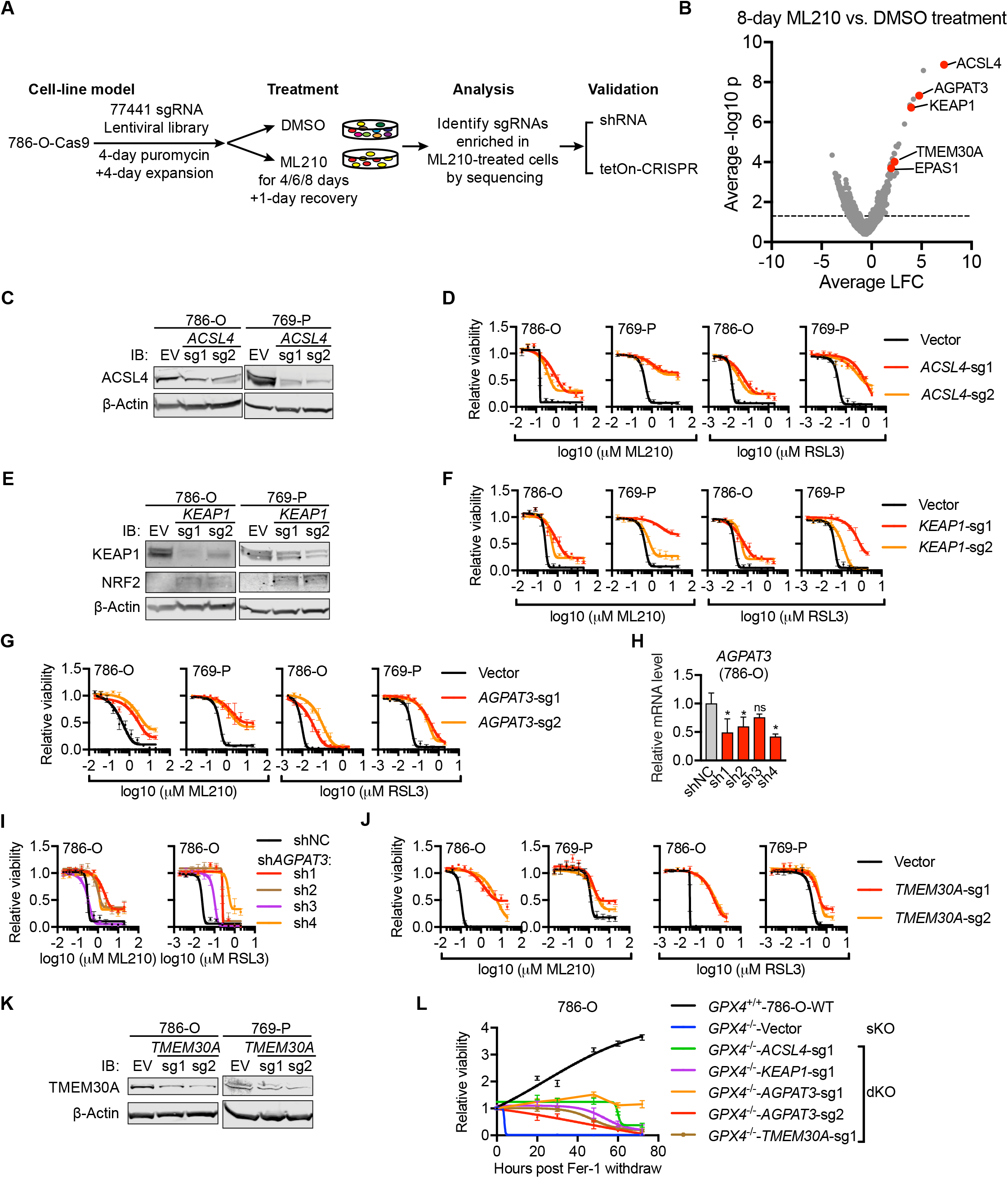
Genome-wide CRISPR resistance screen identifies mediators of ferroptosis susceptibility in ccRCC cells. **A.** Experimental scheme describing the genome-wide CRISPR resistance screening to identify mediators of ML210 sensitivity in 786-O cells. 786-O cells were transduced with Cas9 expression, infected with a genome-wide lentiviral sgRNA library, selected with puromycin, and expanded for treatment with DMSO or ML210 for 4, 6, or 8 days. Forty million cells from the DMSO-treated conditions (minimum representation number per sgRNA> 500) were used as control samples, whereas cells that survived ML210-treatment were grown in drug-free media for one day before harvest. Genomic DNA was extracted from cell pellets and sgRNA barcode abundances were analyzed by sequencing. Top genes with sgRNAs enriched in ML210-treated cells were further validated. **B.** Volcano plot highlighting (red) the top enriched CRISPR hits comparing 786-O cells treated with ML210 or DMSO for 8 days. LFC, log2 fold change (ML210/DMSO). **C.** Immunoblot analysis of ACSL4 protein levels in vector (EV) and ACSL4-targeting sgRNA-expressing 786-O-Cas9 and 769-P-Cas9 cells. β-Actin was used as a loading control. **D.** Viability curves for WT (Vector) and ACSL4-targeting sgRNA-expressing 786-O or 769-P cells under indicated concentrations of ML210 or RSL3. **E.** Immunoblot analysis of KEAP1 and NRF2 *(NFE2L2)* protein levels in WT (Vector) or KEAP1-targeting sgRNA-expressing 786-O-Cas9 and 769-P-Cas9 cells. β-Actin was used as a loading control. **F.** Viability curves for WT (Vector) and KEAP1-targeting sgRNA-expressing 786-O-Cas9 or 769-P-Cas9 cells under indicated concentrations of ML210 or RSL3. **G.** Viability curves for WT (Vector) and AGPAT3-targeting sgRNA-expressing 786-O-Cas9 or 769-P-Cas9 cells under indicated concentrations of ML210 or RSL3. **H.** qRT-PCR analysis of relative *AGPAT3* mRNA expression in 786-O cells expressing shNC or AGPAT3-targeting shRNAs. *B2M* was used as a loading control. Each condition has three biological replicates. Error bars represent ±S.D. Students’ t-test was performed between each shRNA and shNC. *, p<0.05. **I.** Viability curves for shNC or AGPAT3-targeting shRNA-expressing 786-O cells under indicated concentrations of ML210 or RSL3. **J.** Viability curves for WT (Vector) and TMEM30A-sgRNA expressing 786-O-Cas9 or 769-P-Cas9 cells under indicated concentrations of ML210 or RSL3. **K.** Immunoblot analysis of TMEM30A protein levels in WT (EV) and two TMEM30A-targeting sgRNA-expressing 786-O-Cas9 and 769-P-Cas9 cells. β-Actin is used as a loading control. **L.** Viability time-course of the GPX4^+/+^-786-O-WT, *GPX4^−/−^* -Vector (single knockout, sKO), and double knockout (dKO) lines including *GPX4^−/−^-ACSL4-sg1*, *GPX4^−/−^-KEAP* 1-sg1, *GPX4^−/−^-AGPAT3-sg1*, *GPX4^−/−^-AGPAT3-sg2*, and *GPX4^−/−^-TMEM30A-sg*1 cells post Fer-1 withdrawal. Viability is relative to the 0-h condition before Fer-1 withdrawal. Representative plot of experiments repeated three times is presented. For viability assays in panel **D,F,G,I** and **J**, viability under each condition is relative to that of the respective DMSO-treated condition. Each data point has four biological replicates, and error bars represent ±S.D. Representative plot of experiments repeated three times is presented. See also Figure S2.

Notably, our screening also highlighted genes involved in phospholipid regulation including *AGPAT3* and *TMEM30A* (**Figures 2B and S2E**). AGPAT3 (also known as lysophosphatidic acid acyltransferase 3, or LPAAT3) selectively incorporates arachidonic acid (C20:4) and docosahexaenoic acid (C22:6) as acyl-chain donors to lysophosphatidic acids for phosphatidic acid synthesis [58–61]. Depleting AGPAT3 by CRISPR conferred significant resistance to GPX4 inhibition in ccRCC cells (**Figure 2G**). Lacking suitable AGPAT3 antibodies, we verified the effective reduction in *AGPAT3* mRNA levels in AGPAT3-targeting shRNA-expressing cells, which exhibited resistance to GPX4 inhibition (**Figures 2H-I and S2F-G**). These analyses support AGPAT3 as a novel mediator of ferroptosis susceptibility in ccRCC. *TMEM30A* (also known as CDC50A) encodes the β-subunit of the P4-ATPase-associated phospholipid flippase, an enzymatic complex mediating the translocation of phospholipids, particularly phosphatidylethanolamines (PEs), to the inner membrane leaflet and maintaining membrane directionality [62, 63]. TMEM30A-depletion diminished the sensitivity to GPX4 inhibitors in ccRCC cells (**Figures 2J-K**), suggesting that phospholipids on the two leaflets of cellular membranes contribute asymmetrically to ferroptosis.

We then generated double knockouts (dKO) of *GPX4* and *AGPAT3, ACSL4, KEAP1,* or *TMEM30A,* respectively, from *GPX4^−/−^* (single knockout, sKO) cells in the presence of Fer-1 and assessed their viability after Fer-1 withdrawal. All dKO lines exhibited significantly higher survival rates than sKO cells over the tested period (**Figure 2L**), further supporting these genes as positive mediators of ferroptosis susceptibility in ccRCC cells. Together, our screening confirmed known regulators and suggests novel mediators of ferroptosis susceptibility, illuminating a concerted molecular network underlying the GPX4-dependent cell state in ccRCC.

### HIF-2*α* enriches polyunsaturated fatty acyl-lipids and drives GPX4 dependency in ccRCC

We posited that the lineage-specific vulnerability to ferroptosis in ccRCC might be due to tissue-specific expression/activation of the top mediators. While *AGPAT3* and *TMEM30A* exhibit notable enrichment in ccRCC tumors than non-ccRCC cancers in TCGA (**Figures S3A**), our screening highlighted the ccRCC oncogenic driver HIF-2α *(EPAS1)* as a top hit (**Figures 2B and S2E**). HIF-2α is activated in ccRCC tumors but not normal renal tissues [22]; and TCGA analysis revealed that *EPAS1* mRNA is also enriched in ccRCC tumors compared with other cancer types, likely reflecting that *EPAS1* expression is itself positively regulated by HIF-2α activity (**Figures S3A-B and cf. Figure 4A**). These insights imply a new role for the oncogenic HIF-2α pathway in creating a vulnerability to GPX4 inhibition-induced ferroptosis in ccRCC.

We then sought to determine whether HIF-2α is necessary and sufficient for this role. HIF-2α protein is expressed in ccRCC cells 786-O, 769-P, and OS-RC2, but not RTCC line BFTC909 (**Figure S3C**). CRISPR-mediated *EPAS1*-depletion verified HIF-2α as a mediator of ferroptosis susceptibility in ccRCC cells (**Figures 3A-B**). To verify the results from CRISPR-editing in bulk cells, we generated four biallelic *EPAS1^−/−^* single-cell clones. All four clones exhibited significantly reduced expression of known HIF-2α target genes including *CCND1, PLIN2, VEGFA, NDRG1,* and *SLC2A1,* and diminished sensitivity to GPX4 inhibitors (**Figures 3C and S3D-E**). Importantly, exogenous HIF-2α-GFP expression in *EPAS1^−/−^* clones restored sensitivity to GPX4 inhibition (**Figures 3D and S3F**), supporting the specificity of the knockout effect. Additionally, HIF-2α knockout blocked GPX4-depletion-triggered ferroptosis in 786-O cells (**Figure 3E**). The genetic requirement of HIF-2α for ferroptosis susceptibility in 786-O cells is further supported by assessments of EPAS1-shRNA cells (**Figures 3F and S3G-I**). Moreover, HIF-2α-antagonist compound 2 (C2) treatment significantly reduced HIF-2α target gene expression and mitigated GPX4 inhibition-induced ferroptosis in 786-O cells (**Figures S3J-M**) [64]. Notably, cancer cells with *VHL* mutations exhibit greater dependency on GPX4 than *VHL* wildtype cells in a pan-cancer DepMap analysis (**Figure S3N**). Finally, transiently overexpressing HIF prolyl hydroxylase (PHD)-resistant, constitutively active HIF-2α^P405A/P531A^ and HIF-2α^P405A/P531A/N847A^ variants in HK-2 cells increased sensitivity to GPX4 inhibition (**Figure S3O**) [65]. Thus, activated HIF-2α is necessary and sufficient for driving GPX4 dependency in ccRCC cells.

**Figure 3.**
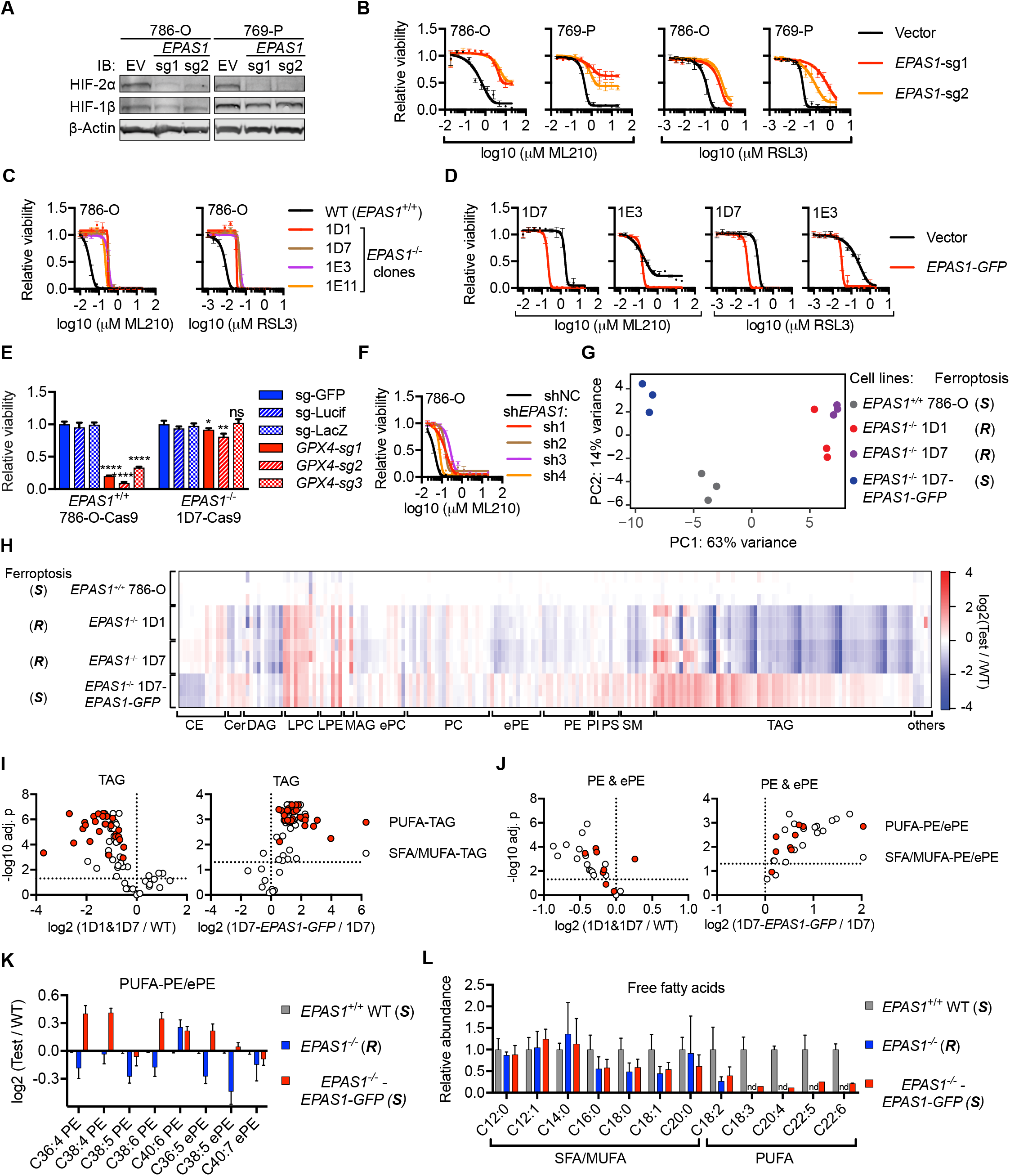
HIF-2α activation drives ferroptosis susceptibility and enriches polyunsaturated fatty acyl (PUFA)-lipids in ccRCC cells. **A.** Immunoblot showing the HIF-2α and HIF-1β protein levels in wildtype (EV) or *EPAS1*-targeting sgRNA-expressing 786-O-Cas9 and 769-P-Cas9 cells. β-Actin is used as a loading control. **B.** Viability curves of wildtype (Vector) or EPAS1-targeting sgRNA-expressing 786-O-Cas9 or 769-P-Cas9 cells treated with indicated concentrations of ML210 or RSL3. **C.** Viability curves for WT 786-O or *EPAS1^−/−^* clones treated with indicated concentrations of ML210 or RSL3. **D.** Viability curves for *EPAS1^−/−^* 786-O single-cell clones 1D7 and 1E3 expressing vector or EPAS1-GFP, then treated with indicated concentrations of ML210 or RSL3. **E.** Relative viability of EPAS1^+/+^786-O-Cas9 and *EPAS1^−/−^* 1D7-Cas9 cells transduced with control (sg-GFP, *sg-Firefly Luciferase*, or sg-LacZ) or GPX4-targeting sgRNAs at day 7 post-infection. Each data point has four biological replicates, and error bars represent ±S.D. Student’s t-test was performed between each of GPX4-targeting sgRNAs and the sg-GFP condition. *, p<0.05, **, p<0.01, ****, p<0.0001, ns, not significant. **F.** Viability curves for 786-O cells expressing shNC or EPAS1-targeting shRNAs treated with indicated concentrations of ML210. **G.** Principal component plot of the lipidomic profiles for the indicated cell lines. (S), ferroptosis-sensitive; *(R),* ferroptosis-resistant. **H.** Heatmap representing the relative lipid abundances in the indicated cell lines. The abundance of each lipid species is normalized to the mean of that in the *EPAS1^+/+^* 786-O WT cells and then log2 transformed. The lipids are grouped by classes, and within each class, the lipid species are ordered first with increasing carbon number, then with increasing unsaturation levels. Abbreviations: CE, cholesterol ester; Cer, ceramide; MAG, monoacylglycerol; DAG, diacylglycerol; TAG, triacylglycerol; LPC, lysophosphatidylcholine; LPE, lysophosphatidylethanolamine; PC, phosphatidylcholine; PE, phosphatidylethanolamine; ePC, (vinyl ether-linked) PC-plasmalogen; ePE, (vinyl ether-linked) PE-plasmalogen; PI, phosphatidylinositol; PS, phosphatidylserine; SM, sphingomyelin. Blue: down-regulated relative to the WT cells, red: up-regulated relative to the WT cells. The “wave”-like pattern in the TAG class corresponds to the more significant losses in the polyunsaturated fatty acyl (PUFA)-TAGs than saturated/monounsaturated fatty acyl (SFA/MUFA)-TAGs in response to HIF-2α-depletion. **I.** Volcano plot showing changes in TAGs grouped as PUFA-TAGs (red fill) and SFA/MUTA-TAGs (white fill) between the indicated cell lines. **J.** Volcano plot showing changes in PE and ePE lipids grouped as PUFA-PE/ePEs (red fill) and SFA/MUFA-PE/ePEs (white fill) between the indicated cell lines. **K.** Bar graph representing the relative abundances of the indicated PUFA-PE/ePE lipids in the labeled groups. Log2 fold changes relative to 786-O WT cells are presented for each condition. Error bars represent ±S.D. **L.** Bar graph representing the relative abundances of the indicated free fatty acids in the labeled groups. nd, not-detectable under the experimental condition. Three biological replicates were included for each condition and error bars represent ±S.D. For viability assays in panel **B,C,D** and **F**, viability under each condition is relative to that of the respective DMSO-treated condition. Each data point has four biological replicates, and error bars represent ±S.D. Representative plot of experiments repeated three times is presented. See also Figure S3.

Ferroptosis is executed by peroxidized membrane phospholipids, particularly PEs that contain polyunsaturated fatty acyl (PUFA)-chains such as arachidonic acid (C20:4) and docosahexaenoic acid (C22:6) [66, 67], and our CRISPR screen highlighted the broad involvement of lipid metabolism genes in ferroptosis susceptibility. We thus hypothesized that HIF-2α might induce ferroptosis sensitization by altering the lipid profile in ccRCC cells. To assess how HIF-2α activation changes lipid metabolism, we performed lipidomic profiling in *EPAS1^+/+^, EPAS1^−/−^* and HIF-2α-GFP rescue 786-O cells (**Figure S3P**). We identified 198 annotated lipids, about 1/3 of which are triacylglycerols (TAG) (**Figure S3Q**), stressing the active lipid deposition in ccRCC cells. HIF-2α-depletion induced a profound shift in the lipidome of 786-O cells, with significant loss of TAGs and phospholipids; and most of these alterations were reverted by HIF-2α-GFP overexpression (**Figures 3G-H and S3R**). Importantly, among all TAGs, PUFA-TAGs exhibited the most significant reduction in response to HIF-2α-depletion compared with saturated/monounsaturated fatty acyl (SFA/MUFA)-TAGs (**Figures 3H-I**). Moreover, levels of most PEs and vinyl ether-linked PE-plasmalogens (ePEs), including PUFA-PE/ePEs, were also significantly reduced in *EPAS1^−/−^* cells, and were then restored by HIF-2α-GFP expression (**Figures 3H,J-K**). Finally, we measured free fatty acid levels in these cells and revealed that free PUFA levels were strongly dependent on HIF-2α activity, whereas free SFA/MUFA levels were less affected by HIF-2α status (**Figure 3L**). These data suggest that HIF-2α promotes the selective enrichment of PUFAs, and stimulates the incorporation of PUFAs into glycerolipids (TAGs and phospholipids) in ccRCC (**Figure S3S**).

### HIF-2α activates *HILPDA* and *G0S2* expression to induce ferroptosis susceptibility

We next sought to determine the downstream mediators of HIF-2α’s ferroptosis-sensitization activity. We first identified HIF-2α-dependent genes and then re-expressed these genes individually in ferroptosis-resistant *EPAS1^−/−^* cells to identify genes that can re-sensitize *EPAS1^−/−^* cells to ferroptosis (**Figure S4A**). RNA-Seq analysis comparing 786-O WT versus *EPAS1^−/−^* cells identified 149 down-and 76 up-regulated genes in response to HIF-2α-depletion, where the top down-regulated genes included *EPAS1* itself and known HIF-2α targets, including *NDRG1, PLIN2, VEGFA* and *SLC2A1* (**Figure 4A**). Notably, at least 11 lipid metabolism genes also exhibited HIF-2α-dependent expression, including *LPCAT1,* a rate-limiting enzyme for phosphatidylcholine (PC) synthesis [68, 69] (**Figure S4B**). HIF-2α-suppressed genes included *CPT1A* and *PPARGC1A* (PGC1 α), each of which promote the mitochondrial translocation and β-oxidation of fatty acids (**Figures 4A and S4B**) [8, 9, 24, 70, 71]. These results verify HIF-2α’s role in suppressing fatty acid β-oxidation and promoting lipid synthesis and deposition [9, 72]. Treatment with HIF-2α antagonist C2 largely recapitulated these gene-expression changes (**Figures 4A and S4B**).

**Figure 4.**
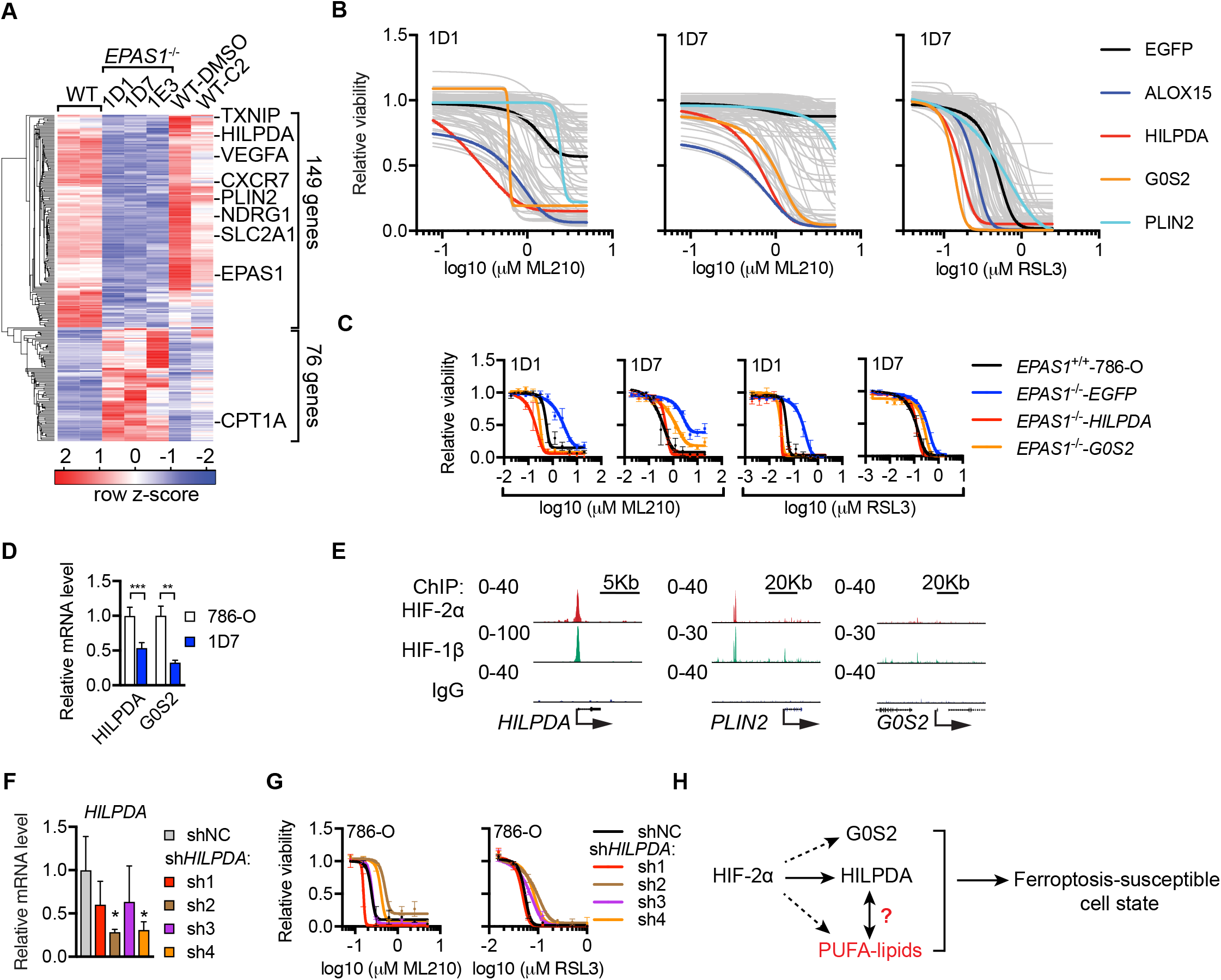
HIF-2α activates *HILPDA* / *G0S2* expression to drive ferroptosis susceptibility in ccRCC cells. **A.** Heatmap showing the RNA-Seq analysis of WT 786-O cells, three *EPAS^−/−^* clones (1D1, 1D7 and 1E3), and DMSO-treated and C2-treated 786-O WT samples. Differential gene expression analysis was performed between WT and *EPAS^−/−^* cells, genes that exhibit fold change >2 or < 0.5 with adjusted p values < 0.05 were presented. Known HIF-2α-responsive genes are highlighted. **B.** Viability curves of cDNA screening results in *EPAS^−/−^* clones treated with a 7-point, 2-fold dilution series of ML210 or RSL3. Viability is relative to that of each cell line under zero ML210 or RSL3 treatment, respectively. *ALOX15* overexpression was used as a positive control. EGFP-expressing *EPAS^−/−^* clones were used as negative controls. Each data point has four biological replicates. Error bars are omitted for visual clarity. **C.** Viability curves of *HILPDA, G0S2,* or EGFP-overexpressing *EPAS^−/−^* 1D1 and 1D7 cells treated with ML210 or RSL3 for 48h. Viability is relative to the respective DMSO-treated conditions. Representative plot of experiments repeated three times. Each data point has four biological replicates, and error bars represent ±S.D. **D.** qRT-PCR analysis of mRNA levels of *HILPDA* and *G0S2* in 786-O-WT and *EPAS1^−/−^* 1D7 cells. Student’s T-test, **, p<0.01, ***, p<0.001. *B2M* was used as an internal control. Representative plot of experiments repeated three times. Each condition includes three biological replicates and error bars represent ±S.D. **E.** Gene-track view for the ChIP-Seq analysis of HIF-2α and HIF-1β in 786-O cells (GSE34871) at the *HILPDA, PLIN2,* and *G0S2* loci. Numbers to the left of each track indicate reads per million reads (RPM). **F.** qRT-PCR analysis of *HILPDA* mRNA levels in 786-O cells expressing shNC or *shHILPDAs. B2M* was used as an internal control. Each condition includes three biological replicates. Error bars represent ±S.D. Student’s T-test. *, p < 0.05. **G.** Viability curves for 786-O cells expressing shNC or *shHILPDAs* under indicated concentrations of ML210 or RSL3 for 48h. Viability is relative to the DMSO-treated conditions. Representative plot of experiments repeated three times. Error bars represent ±S.D. **H.** Scheme summarizing the downstream mediators of HIF-2α for ferroptosis susceptibility. See also Figure S4.

We were able to collect cDNAs for 77 of the 149 HIF-2α-activated genes, including 9 of the 11 lipid metabolism genes. *EPAS1^−/−^* cells expressing each cDNA were assessed for their sensitivity to GPX4 inhibition (**Figure 4B**). As a positive control, we included *ALOX15* (15-LOX), a lipoxygenase that promotes ferroptosis by accelerating lipid peroxidation [34] (**Figures 4B and S4C**). Importantly, *EPAS1^−/−^* cells that overexpressed hypoxia-inducible lipid droplet-associated protein *(HILPDA)* or G0/G1 Switch 2 (G0S2) were significantly more sensitive to GPX4 inhibition than cells expressing most other cDNAs (**Figures 4B-C and S4C-D**). HILPDA and G0S2 share a homologous PNPLA-binding motif and both can act as co-inhibitor of adipose triglyceride lipase (ATGL, encoded by *PNPLA2)* [73, 74], the rate-limiting enzyme for TAG hydrolysis [75] (**Figure S4E**). Notably, overexpressing another HIF-2α-regulated, ccRCC-enriched, and lipid droplet-associated protein perilipin2 *(PLIN2)* [9, 76] did not alter GPX4 inhibitor sensitivity in the same cells (**Figure 4B and S4B-C**).

*HILPDA* and *G0S2* expression are both induced by hypoxia [77, 78]. *HILPDA* was shown to be highly expressed in RCCs but not in normal renal tissue [79], and our TCGA analysis highlighted ccRCC tumors as the highest HILPDA-expressing cancers in average (**Figure S4F**). *G0S2* is expressed in most cancer types, with ccRCC among the highest G0S2-expressing cancers (**Figure S4G**). qRT-PCR analysis verified HIF-2α-dependent *HILPDA* and *G0S2* expression in 786-O cells (**Figure 4D**). Notably, previously reported ChIP-Seq data revealed strong HIF-2α/HIF-1β binding to *HILPDA* and *PLIN2* promoters in 786-O cells [80]; however, no significant HIF-2α binding has been identified near the *G0S2* locus (**Figure 4E**) [80]. These results indicate that *HILPDA* is a direct target of HIF-2α, whereas *G0S2* expression is regulated by HIF-2α via indirect mechanisms. shRNA-mediated knockdown of endogenous *HILPDA* partially, albeit significantly, decreased GPX4 inhibitor sensitivity in 786-O cells (**Figures 4F-G**). Nonetheless, G0S2-shRNA did not induce similar effects (**Figures S4H-J**), potentially due to functional compensation from HILPDA. Considering that exogenous HILPDA induced comparable sensitization effects as G0S2 while lower HILPDA protein levels were detected in the overexpression system (**Figure S4C**), we concluded that HILPDA is a leading mediator of HIF-2α’s activity in driving ferroptosis susceptibility, whereas G0S2 plays a facilitatory role (**Figure 4H**).

### HIF-2α–HILPDA axis drives a polyunsaturated fatty acyl-lipid-enriched state in ccRCC

We next characterized the mechanisms by which HILPDA and G0S2 mediate ferroptosis susceptibility in ccRCC. HILPDA and G0S2 proteins are localized on lipid droplets (LDs), and a predominant consequence of *G0S2* expression in adipocytes is diminished lipolysis and aberrant accumulation of LDs [81–83]. Notably, increased LD abundances were observed in *G0S2-* or *PLIN2-* but not HILPDA-overexpressing *EPAS1^−/−^* cells (**Figure S5A**), suggesting that nascent LD accumulation is neither necessary nor sufficient in mediating ferroptosis susceptibility in ccRCC; instead, specific contents in LDs may play a more direct role in ferroptosis. This result also suggests that HILPDA drives ferroptosis susceptibility through other mechanisms rather than regulating LD abundances. By lipidomic profiling, we revealed that HILPDA-overexpression largely restored the lipidomic changes induced by HIF-2α-depletion in *EPAS1^−/−^* cells, while *G0S2-* overexpression induced similar trends with notable distinctions (**Figures 5A-B and S5B-C**).

**Figure 5.**
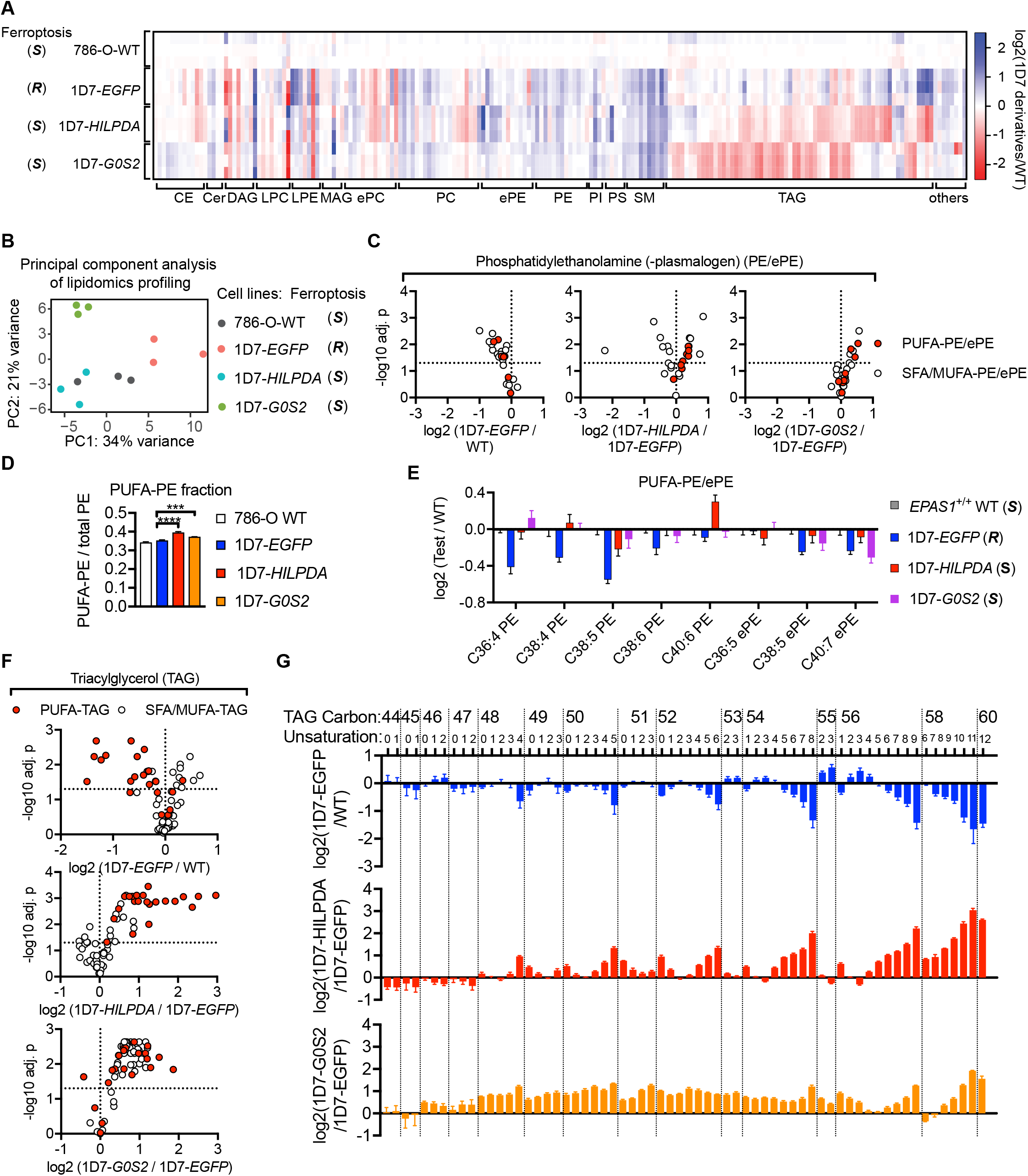
The HIF-2α-HILPDA axis drives the selective enrichment of PUFA-TAGs and PUFA-phospholipids. **A.** Heatmap representing the relative lipid abundances in the indicated cell lines. The lipid ratios between 1D7 derivatives and 786-O WT cells were log2 transformed and plotted. Lipid class abbreviations are the same as in Figure 3F. (S), ferroptosis-sensitive; (R), ferroptosis-resistant. **B.** Principal component plot of lipidomic profiles for the indicated cell lines. **C.** Volcano plots showing the changes in each PE/ePE species grouped as PUFA-PE/ePE (red fill) and SFA/MUFA-PE/ePEs (white fill) between the indicated cell lines. **D.** The ratio between PUFA-PE/ePE and total PE/ePE in the indicated cell lines. **E.** Bar graph showing the relative abundances of the PUFA-PE/ePEs in the conditions tested. The lipid ratios between 1D7 derivatives and 786-O WT cells were log2 transformed and plotted. Each condition includes three biological replicates. Error bars represent ±S.D. **F.** Volcano plots showing the changes in each TAG species grouped as PUFA-TAGs (red fill) and SFA/MUTA-TAGs (white fill) between the indicated cell lines. **G.** Bar graph showing the relative abundances of each TAG species between the cell lines indicated. The lipid ratios between 1D7 derivatives and 786-O WT cells were log2 transformed and plotted. Error bars represent ±S.D. See also Figure S5.

Remarkably, HILPDA selectively restored the levels of most PUFA-PE/ePEs, including C36:4, C38:4 and C40:6 PEs as well as C38:5 and C40:7 ePEs (**Figures 5C-E**). G0S2, instead, induced a global up-regulation of the PE/ePEs class (**Figures 5C-E**). Similarly, *EPAS1^−/−^-HILPDA* cells exhibited substantial enrichment of PUFA-TAGs but not SFA/MUFA-TAGs, whereas *EPAS1^−/−^ G0S2* cells non-selectively enriched most TAGs (**Figures 5F-G**). These data suggest that HIF-2α activates *HILPDA* expression to selectively enrich PUFA-lipids, while in parallel inducing G0S2 to generally enhance lipid synthesis and deposition. Both activities together create a robust ferroptosis-susceptible state in ccRCC cells (**Figure S5D**).

### ACSL4 and AGPAT3 selectively incorporates polyunsaturated fatty acids to phospholipids

Our genetic and lipidomic characterizations along the HIF-2α–HILPDA axis highlight a strong correlation between ferroptosis susceptibility and the abundances of PUFA-lipids, including TAGs and phospholipids. While TAGs are aberrantly accumulated in ccRCC cells, they are stored in lipid droplets that are separated from membrane-anchored lipoxygenases (LOXs). Most phospholipids, however, are present on cellular membranes that are readily accessible by LOXs. Nonetheless, how PUFAs are selectively incorporated into phospholipids in ccRCC is poorly understood. Among the top ferroptosis-susceptibility mediators identified in our CRISPR screen, ACSL4 and AGPAT3 exhibit considerable selectivity towards utilizing PUFAs for synthesizing fatty acyl-CoAs and phosphatidic acids (PAs), respectively [53, 58, 59] (**Figure 6A**). PUFA-PAs can be directed towards the synthesis of either TAGs by activities of diglyceride acyltransferases (DGAT1/2) or phospholipids via the CDP-DAG pathway (**Figure 6A**) [84, 85]. Therefore, ACSL4 and AGPAT3 are plausible candidates for mediating PUFA-phospholipid synthesis. Lipidomic profiling revealed that ACSL4-or AGPAT3-knockout led to marked down-regulation of PUFA-PE/ePEs and up-regulation of PUFA-TAGs, whereas SFA/MUFA-lipids (TAGs and phospholipids) are largely insensitive to ACSL4/AGPAT3-depletion (**Figures 6B-E and S6A-D**). These results indicate that ACSL4 and AGPAT3 promote the selective synthesis of PUFA-phospholipids rather than TAGs in ccRCC, and thus modulate the sensitivity to ferroptosis (**Figure S6E**). Notably, the PUFA-TAG species enriched upon ACSL4-or AGPAT3-depletion largely overlapped with the HIF-2α-HILPDA axis-dependent TAGs (**Figure 6F**). This result implies a crosstalk between the HIF-2α-HILPDA and ACSL4/AGPAT3 pathways, potentially through controlling the abundance of free PUFAs available for TAG or phospholipid synthesis (**Figure S6E**). Together, the overall lipogenic, PUFA-lipid-enriched cell state associated with high ferroptosis susceptibility in ccRCC is concertedly regulated by the HIF-2α-HILPDA and ACSL4/AGPAT3 pathways.

**Figure 6.**
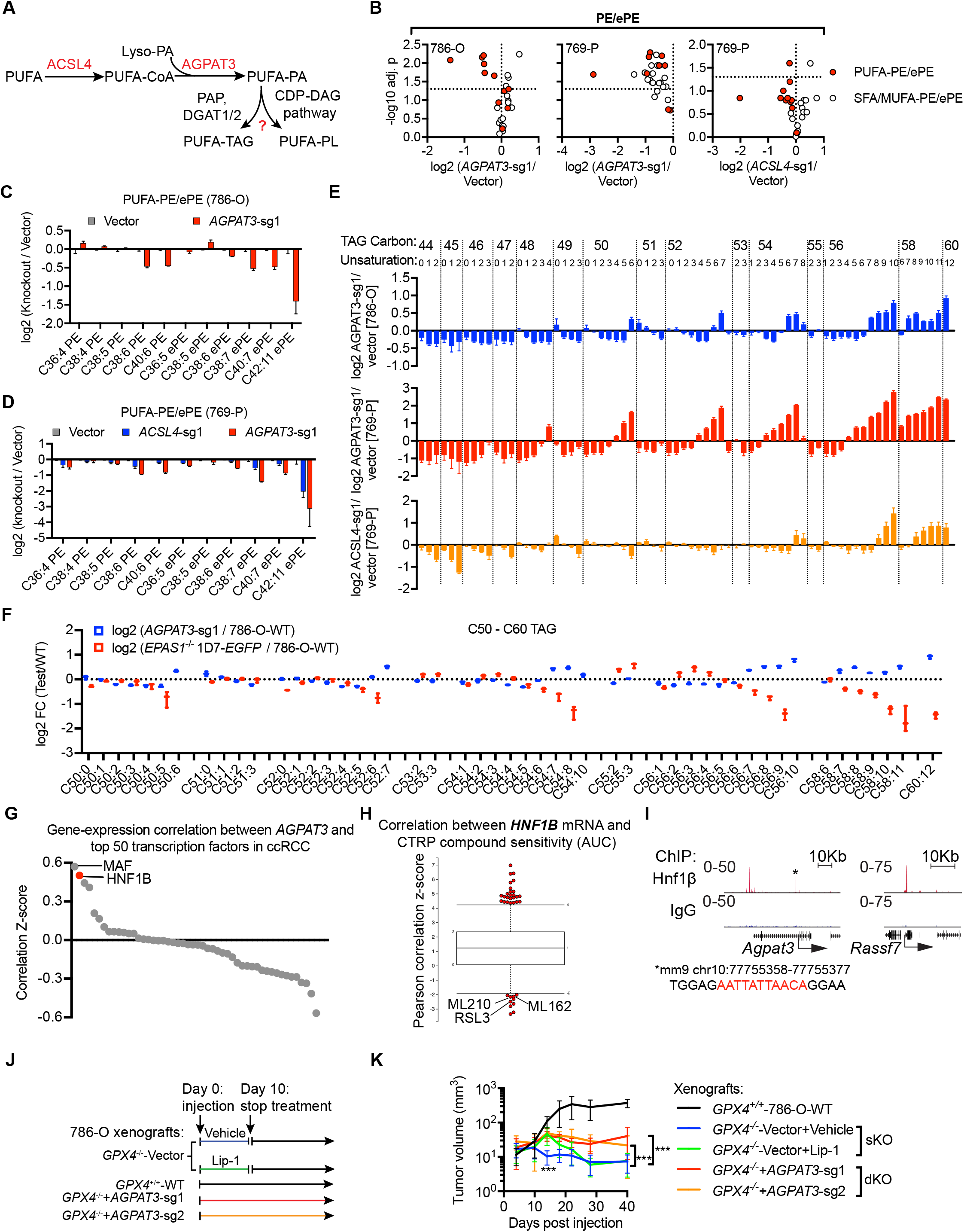
HNF-1β-driven AGPAT3 selectively synthesizes PUFA-phospholipids and primes the renal lineage for GPX4 dependency. **A.** Scheme summarizing the role of ACSL4 and AGPAT3 in PUFA-lipid synthesis, and the potential flux of PUFA-phosphatidic acids (PUFA-PA) towards either PUFA-TAG or PUFA-phospholipid synthesis through distinct enzymatic pathways. **B.** Volcano plots showing changes in each PE/ePE species grouped as PUFA-PE/ePEs and SFA/MUFA-PE/ePEs between the indicated cell lines. Each condition includes three biological replicates. **C.** Bar graph showing relative abundances of the PUFA-PE/ePE lipids in 786-O cells expressing control vector or *AGPAT3-sg1*. Log2 fold changes between the knockout test cells and the 786-O-Cas9 WT cells were plotted. Error bars represent ±S.D. **D.** Bar graph showing the relative abundances of the PUFA-PE/ePE lipids in 769-P cells expressing control vector, ACSL4-sg1, or AGPAT3-sg1. **E.** Bar graph showing the relative abundances of each TAG species in the cell lines indicated. **F.** Relative changes in the abundances of C50-C60 TAGs in *AGPAT3-sg1* or 1 *D7-EPAS1^−/−^* cells versus 786-O-Cas9 WT cells. Error bars represent ±S.D. **G.** Gene-expression correlation analysis between *AGPAT3* and the top 50 RCC-enriched transcription factors including *HNF1B* (red) in the RNA-Seq dataset of ccRCC tumor samples (N=535) from the TCGA database. Correlation z-scores are plotted using Fisher’s z-transformation on raw correlation coefficients. **H.** Compound sensitivity-gene-expression correlation analysis in CTRP, highlighting that *HNF1B* expression is strongly correlated with sensitivity to GPX4 inhibitors ML210, RSL3, and ML162. Negative z-score means high expression is correlated with high sensitivity to compound, and vice versa. **I.** ChIP-Seq analysis of (data from GSE71250) Hnf-1β binding sites at the *Agpat3* and *Rassf7* loci in mouse renal mIMCD cells. Track height, reads per million reads (RPM). Bottom: highlight of Hnf-1β consensus motif at the *Agpat3* promoter-binding peak center. **J.** Scheme summarizing xenograft experiments with the indicated cell lines. *GPX4^+/+^* 786-O-WT, *GPX4^−/−^* (sKO), or *GPX4^−/−^* cells expressing AGPAT3-sgRNA1 and sgRNA2 (dKOs) were subcutaneously implanted into immunocompromised nude mice. Matrigel containing **J.** 5μM Lip-1 was used to assist the initial implantation of the cancer cells. Additionally, mice bearing *GPX4^−/−^* (sKO) tumors were separated into two groups, with one group treated with vehicle and the other group treated with Lip-1 daily for 10 days. **K.** Time course of tumor volume measurements of indicated xenografts (five mice per group, two tumors per mouse). Error bars represent ±S.D. Student’s t-test is performed between the volumes of vehicle and Lip-1 treated *GPX4^−/−^* (sKO) tumors at day 14, and between vehicle treated *GPX4^−/−^* (sKO) tumors and *GPX4^−/−^-AGPAT3-sg1* or *GPX4^−/−^-AGPAT3-sg2* (dKO) tumors at day 40. ***, p<0.001. See also Figure S6.

**Figure 7.**
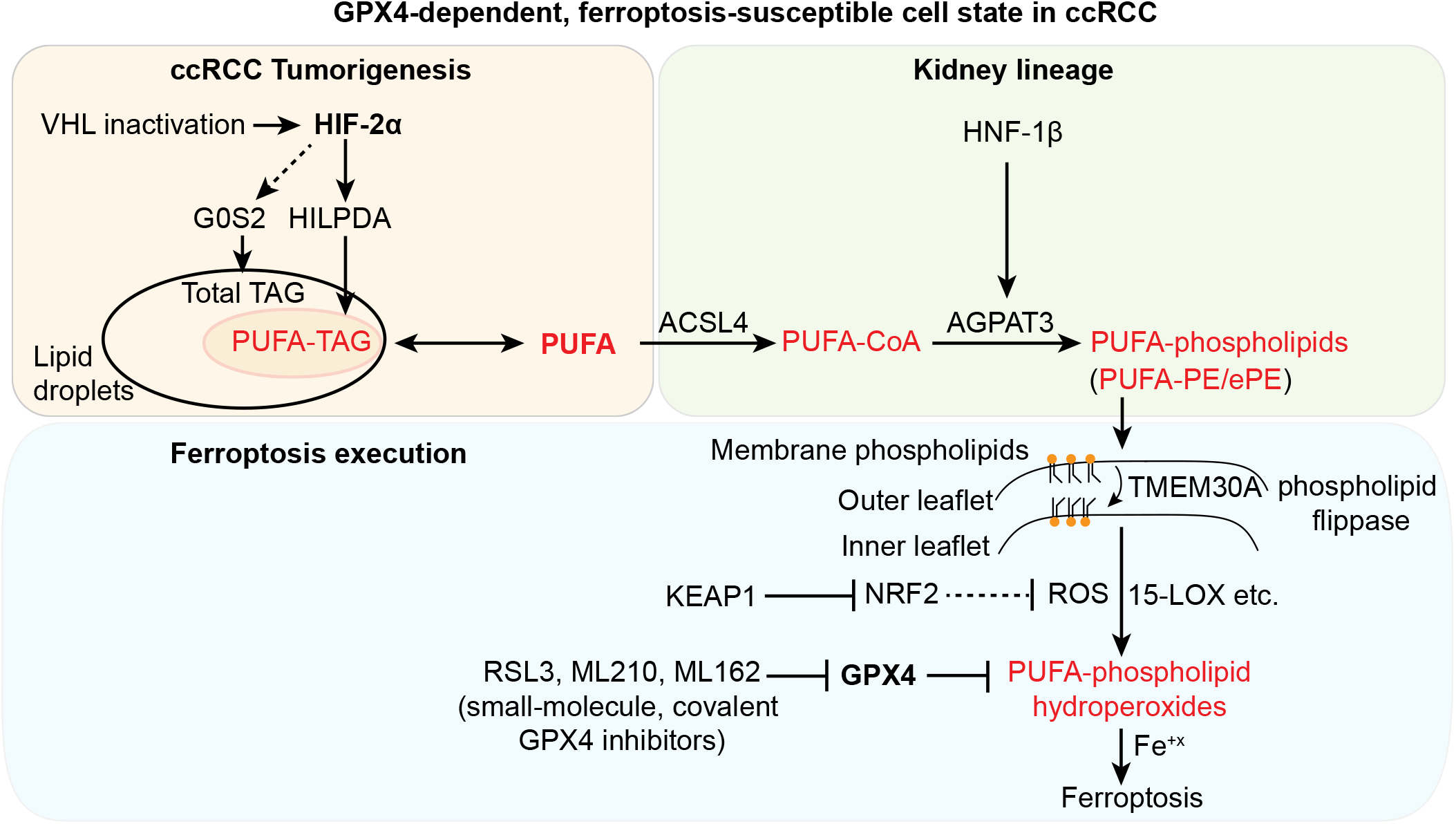
Scheme summarizing the molecular network driving the intrinsic GPX4 dependency and ferroptosis susceptibility in ccRCC. See Discussion. Abbreviations: ccRCC, clear cell renal cell carcinoma; PUFA, polyunsaturated fatty acids, e.g. arachidonic acid (C20:4); TAG, triacylglycerols; PE, phosphatidylethanolamine; ePE, vinyl ether-linked PE-plasmalogens; ROS, reactive oxygen species; 15-LOX, 15-lipoxygenase. Metabolites highlighted in red indicate promoters of ferroptosis susceptibility. Dashed arrow lines represent indirect regulation; double arrow represents bidirectional metabolic flow.

To verify the lipidomic changes associated with ferroptosis susceptibility in clinically relevant settings, we interrogated a previously reported lipidomic profiling dataset comprising 49 ccRCC tumor/normal tissue pairs [86]. All TAGs, including PUFA-TAGs, were up-regulated in tumors compared with normal tissues, consistent with our results and the “clear-cell” hallmark [86]. Importantly, ccRCC tumors exhibited higher levels of PUFA-PE/ePEs and PUFA-PC/ePCs, which were further enriched in high-grade tumors (stage III/IV) when compared with low-grade ones (stage I/II) (**Figures S6F-G**). These results support that human ccRCC tumors are in a PUFA-lipid-enriched state.

### HNF1β regulates *AGPAT3* expression and primes renal cells for ferroptosis

Considering AGPAT3’s significant enrichment in ccRCC tumors and kidney cell lines (**Figure S6H-I**), and its remarkable selectivity in driving PUFA-phospholipid synthesis, we explored the upstream regulators of *AGPAT3* in the renal lineage. By correlating the expression of the top 50 transcription factors (TFs) with *AGPAT3* mRNA levels in the TCGA ccRCC dataset, we found *MAF* and *HNF1B* as the most significant AGPAT3-positively correlated TFs (**Figure 6G**). Notably, *HNF1B* expression is also strongly correlated with high sensitivity to GPX4 inhibitors in CTRP (**Figure 6H**)[15]. HNF-1β is a lineage-specification factor in renal tubule development, and is essential for ccRCC cell survival [45, 87], which may account for the lack of HNF-1β sgRNA representation in our CRISPR screen. Analysis of previously reported Hnf-1β ChIP-Seq data in mouse renal IMCD cells confirmed strong Hnf-1β binding to *Agpat3* promoter and putative enhancer with comparable intensities to the strongest Hnf-1β peak at *Rassf7* promoter in this dataset (**Figure 6I**) [88]. Additionally, an Hnf-1β consensus-binding motif was identified at the *Agpat3* promoter (**Figure 6I**). These results support that *AGPAT3* expression is driven by HNF-1β in renal cells, and that the HNF-1β-AGPAT3 axis primes renal cells for ferroptosis prior to tumorigenesis.

We next assessed AGPAT3’s role in ferroptosis *in vivo* by implanting a *GPX4^−/−^* (sKO) line, two *GPX4/AGPAT3* double knockout (dKO) lines, and WT 786-O-Cas9 cells into immunodeficient mice, with the *GPX4^−/−^* sKO mice being treated with either vehicle or Lip-1 daily for 10 days (**Figure 6J**). In contrast to the rapid regression of *GPX4^−/−^* tumors treated with vehicle, Lip-1 treated tumors attained significantly larger sizes before regressing upon Lip-1 withdrawal (**Figure 6K**), confirming that GPX4-null cells underwent ferroptosis *in vivo.* Importantly, *GPX4/AGPAT3* dKO tumors continued to establish tumor nodules, though at a slower rate than the WT counterparts (**Figure 6K**), suggesting that AGPAT3 is necessary for ferroptosis in ccRCC tumors *in vivo.* Together, the HNF-1β-AGPAT3 axis is a critical determinant of the vulnerability to ferroptosis in ccRCC.

## DISCUSSION

Clear cell renal cell carcinoma (ccRCC), while curable in the localized setting, is incurable upon recurrence which can happen in 30% of all curative-intent nephrectomies. Furthermore, 20-30% of ccRCC patients present with de-novo metastatic disease. The metastatic form of ccRCC kills thousands of patients annually in the US [89], despite recent advances in systemic therapy such as anti-angiogenics and novel checkpoint blockers. Here, we uncover an intrinsic susceptibility to ferroptosis and GPX4 dependency in ccRCC. Through a combination of small-molecule sensitivity profiling, genome-wide CRISPR screening, metabolomic profiling, and functional cDNA screening, we reveal that this vulnerability is rooted in the molecular circuitry of the renal lineage and ccRCC tumorigenesis (**Figure 7**). In ccRCC cells, activated HIF-2α drives *HILPDA* expression to selectively enrich polyunsaturated fatty acyl-triacylglycerols (PUFA-TAGs), while in parallel inducing G0S2 and suppressing fatty acid β-oxidation to up-regulate global lipid synthesis and deposition. These activities lead to increased abundances of free fatty acids, particularly PUFAs, which are selectively incorporated into PUFA-phospholipids by the sequential activities of ACSL4 and AGPAT3. Elevated PUFA-phospholipids, particularly PUFA-PE/ePEs, create a strong vulnerability toward ferroptosis and thus a dependency on GPX4 (**Figure 7**).

While HIF-2α activation is acquired by genetic or epigenetic inactivation of *VHL, AGPAT3* expression is activated by the HNF-1β transcriptional machinery in renal epithelial cells prior to transformation. The HNF-1β-AGPAT3 axis promotes PUFA-phospholipid synthesis and primes renal cells for GPX4 dependency and the potential for ferroptosis following GPX4 inhibition. This pathway especially results in PUFA-PE/ePEs translocating to the inner leaflet of cellular membranes by the actions of the phospholipid flippase TMEM30A, where they are subject to peroxidation catalyzed by membrane-anchored lipoxygenases. Under normal growth conditions, cells counter phospholipid hydroperoxide accumulation and evade ferroptosis through the lipid hydroperoxide-selective activity of GPX4 and more generally the NRF2-driven anti-oxidant program. GPX4 inhibition directly triggers, whereas KEAP1-depletion mitigates ferroptosis in ccRCC cells (**Figure 7**). While ccRCC exhibits broad resistance to apoptosis-inducing agents, its unique vulnerability to ferroptosis offers a new opportunity to conquer this devastating disease.

Our analysis agrees with prior studies showing that ferroptosis is triggered in renal tubule cells by conditional *Gpx4* knockout and folic acid or ischemia/reperfusion-induced acute renal injury [30, 48, 90]. The fact that various cellular insults induce the same cell death mechanism in kidney-originated cells suggest that the renal lineage is primed to undergo ferroptosis. Here we demonstrate a potential mechanism underlying this primed state, involving the HNF-1β–AGPAT3-PUFA-phospholipid axis that is highly active in the renal lineage. Consistent with our data, previous report showed that the renal cortex, where proximal tubules are located, exhibits higher levels of PUFA-lipids than the medulla [91]. Moreover, our study underscores an increase in ferroptosis susceptibility that is paradoxically driven by oncogenic HIF-2α in ccRCC. While ferroptotic cell death was first demonstrated in the context of selective sensitivity in fibroblasts engineered with forced expression of an oncogenic allele of HRAS [47, 92], our work underscores a prominent example of “oncogenesis-associated ferroptosis susceptibility” in human cancers. The involvement of HIF-2α in ferroptosis was indirectly supported by a recent report showing that pVHL restoration reverted the sensitivity to erastin and L-buthionine-S,R-sulfoximine (BSO) in RCC4 cells, although the mechanisms mediating pVHL’s action was not elucidated [29], and non-ferroptotic contributions to death by these two agents are suggested by the partial rescue using ferrostatin-1. Here we report that HIF-2α selectively enriches PUFAs rather than SFA/MUFAs, up-regulates the abundances of PUFA-TAGs as well as PUFA-phospholipids, and induces a GPX4-dependent, ferroptosis-susceptible state in ccRCC cells. Hence, our characterizations provide a conceptual framework for understanding ferroptosis sensitivity in renal tissues at the baseline and the marked increase in this vulnerability as a result of oncogenic transformation (**Figure 7**).

Our observations about the elevated ferroptosis sensitivity in ccRCC cell lines and primary cultures relative to normal renal cells suggest a potential therapeutic window for inducing ferroptosis to treat ccRCC tumors. While most current apoptosis-inducing treatment modalities for kidney cancer are challenged by complications from low response rates and emergences of resistance, ferroptosis-inducing agents potentially offer a new paradigm in the way that kidney cancers can be targeted. Given GPX4’s unique substrate specificity and the lack of functional redundancy from other peroxidases, directly targeting GPX4 to induce ferroptosis is an appealing approach for developing new ccRCC therapies. However, due to the poor bioavailability of current small molecule GPX4 inhibitors, the *in vivo* efficacy of chemical inhibition of GPX4 in ccRCC models remains to be demonstrated. While a recently reported GPX4-targeting strategy suggests that it may be possible to overcome the liabilities associated with conventional GPX4 inhibitors [93], developing novel GPX4 inhibitors with improved pharmacokinetics and pharmacodynamics profile warrants further investigation.

While our functional screening was performed in ccRCC cells, many of the top hits are involved in fundamental aspects of redox homeostasis, lipid metabolism and cancer progression, suggesting that their roles in ferroptosis susceptibility can be extrapolated to other cancers. Though few cell-line models of other kidney cancer subtypes, such as papillary and chromophobe RCCs, are available for dependency analyses, all RCCs express *AGPAT3* and *HNF1B* at high levels and are likely to be partially sensitive to ferroptosis. Renal oncocytomas, which are benign cancers originated from the distal nephron, exhibit alterations of glutathione metabolism indicative of GPX4 dependency (GPX4 uses glutathione as an obligate cofactor). Remarkably, a fumarate hydratase-deficient hereditary leiomyomatosis renal cell carcinoma cell line UOK262 was recently characterized as ferroptosis-susceptible through mechanisms involving fumarate-mediated post-translational modification thus inactivation of GPX4 [94]. In extrarenal cancers, clear-cell ovarian carcinoma cells exhibit elevated levels of HIF-2α, HILPDA, and HNF-1β expression [95–97], and these cells are highly dependent on GPX4 in our CTRP analysis. Other candidates for high ferroptosis susceptibility include pheochromocytoma and paraganglioma (PCPG), two are neuroendocrine malignancies that are frequently associated with mutations in the VHL/HIF pathway and exhibit aberrant cytoplasmic lipid accumulation [98]; and uveal melanoma, which exhibits high *AGPAT3* expression (**Figure S6H**). Though these cancers are not well represented in CTRP or DepMap, they all lack effective therapies and therefore testing their GPX4 dependency merits future investigation. Moreover, though the physiological role of PUFA-TAGs/phospholipids in cancers remain unclear, cancer stem cells in several contexts exhibit increased lipid unsaturation levels [15, 99], pointing towards broader applications of ferroptosis-inducing agents in conquering cancer stemness and metastasis.

In addition to the potential impact on cancer therapy, this study also provides insights into the mechanisms underlying ferroptosis. First, our genetic and lipidomic analyses clarify the contributions of PUFA-phospholipids and PUFA-TAGs, the two major pools of PUFA-lipids, in the execution of ferroptosis. The insensitivity to ferroptosis in ACSL4-or AGPAT3-depleted cells, both of which exhibit increased PUFA-TAGs and decreased PUFA-phospholipids, argues against the possibility that PUFA-TAGs directly trigger ferroptosis and supports phospholipids as the ferroptosis substrate. Instead, a PUFA-TAG and PUFA-phospholipid interconversion model is suggested. While lipid droplets (LDs) in clear-cell carcinomas have long been considered labile organelles that protect cells from lipotoxicity [9, 100], our data imply that PUFA-TAGs in LDs of ccRCC contribute to ferroptosis susceptibility by acting as reservoirs of PUFAs in non-toxic forms and sources for PUFA-phospholipid synthesis. Recently, a mutant allele of *PNPLA3,* the closest homolog of ATGL *(PNPLA2),* was recently reported to selectively convert PUFA-TAGs to PUFA-phospholipids in non-alcoholic fatty liver disease [101]. While *PNPLA3* mRNA levels appear below detection limit in ccRCC cells (data not shown), the mechanisms that mediate PUFA-TAG to PUFA-phospholipid conversion remain to be elucidated.

Secondly, our study emphasizes the potent activity of HILPDA in inducing a ferroptosis-susceptible state downstream of HIF-2α in ccRCC cells. This activity is likely mediated through HILPDA’s previously dismissed selectivity towards enriching PUFA-TAGs/phospholipids over SFA/MUFA-lipids. Although HILPDA and G0S2 were both shown to repress ATGL activity [73, 82, 83], the distinct lipidomic profiles in *HILPDA-,* and G0S2-overexpressing cells characterized here strongly suggest that an ATGL-independent activity of HILPDA is present. This activity, coupled with the remarkable selectivity towards PUFA-lipids, is likely contributing to ferroptosis susceptibility in ccRCC and other contexts. Deciphering the underlying biochemical mechanisms of such activity merits future study. Additionally, our results suggest that HILPDA expression may be used as a biomarker to predict sensitivity to GPX4-targeting agents in patients. Finally, as HIF-2α drives multiple branches of downstream events during ccRCC tumorigenesis, it will be intriguing to examine whether tumors that acquire resistance to HIF-2α inhibitors retain their ferroptosis susceptibility and sensitivity to GPX4 inhibition.

Lastly, the involvement of phospholipid flippase component TMEM30A in ferroptosis highlights the potential asymmetric contributions of the two membrane leaflets and points towards the inner leaflet as more directly involved in creating susceptibility to ferroptosis. A plausible explanation is that PE/ePEs are preferentially translocated to the membrane inner leaflet by phospholipid flippases, thus more readily exposing PUFA chains to intracellular ROS and membrane-associated lipoxygenases or other enzymes involved in lipid peroxidation [102–104]. Nonetheless, the precise mechanisms by which membrane topology affects ferroptosis kinetics remain to be characterized. Notably, the impact on ferroptosis susceptibility by membrane asymmetry is reminiscent of the “flip-flop” model of membrane phosphatidylserines during apoptosis [105, 106], which stresses the remarkable similarities between these two evolutionarily conserved pathways – both involving membrane topology for regulating the dynamics of cell death.

Our CRISPR screen, like resistance screens for many other drugs, sheds light on potential resistance mechanisms to GPX4-targeted therapies. Genetic or epigenetic suppression of the ferroptosis-susceptibility mediators is a plausible mechanism by which tumors could evade ferroptosis. Our study will guide the development of combination therapies (as sequential or frontline strategies) if resistance to GPX4-inhibitors arises. Importantly, because the ferroptosis-susceptible state in ccRCC shares activated HIF-a status and oxidative stress with ischemia/reperfusion-induced kidney injury [30, 34, 107, 108], the ferroptosis-susceptibility mediators identified in this study are potential drug targets to prevent renal cell death. Moreover, the broad tissue expression patterns of these mediators, including *AGPAT3, ACSL4, KEAP1,* and *TMEM30A,* support the likely involvement of these genes in other ferroptosis-related diseases beyond the kidney.

In summary, we identify an oncogenesis-associated ferroptosis susceptibility and GPX4-dependency in ccRCC, delineate the genetic and metabolic basis for this dependency, and illuminate the principles of ferroptotic cell death. These insights are potentially translatable towards novel therapies for ccRCC and other diseases involving ferroptosis.

## ACKNOWLEDGEMENTS

We thank Shubhroz Gill, Olivia Bare, Xi Wang, Xin Jin, Xin Rong and Robert L. Bowman for discussion, and Project Achilles for sharing Cas9-expressing cell lines. We thank Cindy Suk-Yee Hon for project management and organization assistance. We thank the William Kaelin laboratory for sharing the HIF-2α^P405A/P531A^ and HIF-2α^P405A/P531A/N847A^ plasmids and the Eric Jonasch laboratory for the HIF-2α-GFP plasmid. The authors acknowledge funding support for M.J.P. from the NIH NHLBI award T32HL007627, and for V.S.V. from the NCI SPORE P50 CA101942-13 Directors Choice Award. This work was supported by the NCI’s Cancer Target Discovery and Development (CTD^2^) Network (grant number U01CA217848, awarded to S.L.S.), and in part by NIH/NCI DF/HCC Kidney Cancer SPORE P50 CA101942 to S.S. and T.K.C., and by the Trust Family, Loker Pinard and the Michael Brigham Funds for Kidney Cancer Research to T.K.C. S.L.S. is an Investigator at the Howard Hughes Medical Institute.

## AUTHOR CONTRIBUTIONS

Y.Z. and S.L.S. conceived the project, designed the experiments and wrote the manuscript. Y.Z. performed experiments and bioinformatics analysis. A.D. and C.B.C. assisted the metabolomics analyses. J.D. assisted the genetic screening experiments and data analysis. M.J.P. and M.A. assisted the imaging experiments. H.L., V.D., E.S.L., V.S.V., B.K.W. and P.A.C. assisted in bioinformatics analysis and manuscript preparation. W.W. and J.K.E. performed chemical synthesis. Y-Y.T., R.D., S.S., T.K.C. and J.S.B. established the primary RCC cell lines. All authors interpreted data, discussed results and contributed to writing the manuscript.

## CONTACT FOR REAGENT AND RESOURCE SHARING

Further information and requests for resources and reagents should be directed to Yilong Zou or Stuart L. Schreiber.

## SUPPLEMENTARY FIGURE LEGENDS

**Figure S1.**
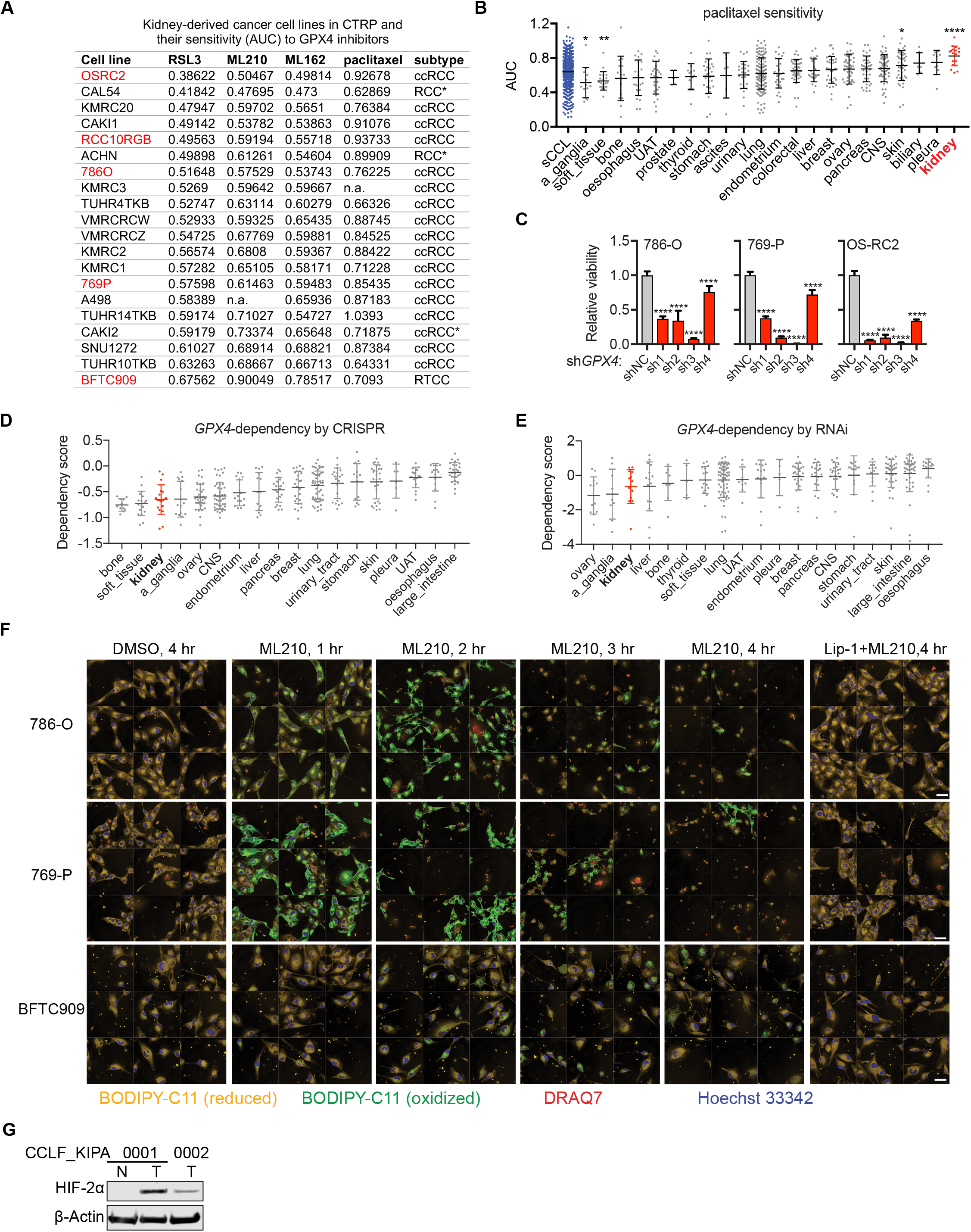
Analysis of Cancer Therapeutics Response Portal (CTRP) data identifies GPX4 as a lineage-specific dependency in ccRCC. Related to Figure 1. **A.** Table summarizing the kidney-derived cancer cell lines characterized in CTRP, their sensitivities to GPX4 inhibitors and paclitaxel by AUC values and the cancer subtype information. * uncertain subtype; n.a., sensitivity value not available. Cell lines marked in red were used for individual validation in **Figure 1F**. **B.** Scatterplot of AUC value distributions for paclitaxel in all solid tumor cancer cell lines (sCCL, blue) or cell lines from each specified tissue-of-origin, including kidney (orange). Larger AUC values indicate lower compound sensitivity, and vice versa. Abbreviations: CNS, central nervous system; UAT, upper aerodigestive tract; a_ganglia, autonomic ganglia. A Mann-Whitney-Wilcoxon test was performed between each tissue and sCCLs from other tissues. Statistical significance is adjusted for multi-test correction. *, p<0.05; **, p<0.01; ****, p< 0.0001. Line and error bars: mean and standard deviation (S.D.). **C.** Relative viability of 786-O, 769-P, and OS-RC2 cells expressing shNC or shGPX4. Cellular viability was measured at 7 days post-lentiviral shRNA infection and normalized to shNC cells. Student’s t-test was performed between each shRNA and the shNC. ****, p<0.0001. Each condition has four biological replicates and error bars represent ±S.D. **D.** Relative sensitivity score (CERES score) of *GPX4* knockout by CRISPR in cancer cell lines from indicated tissue-of-origin including kidney (red) from the Cancer Dependency Map (DepMap) database. Tissue abbreviations are the same as in panel C. **E.** Relative sensitivity score (ATARiS score) of *GPX4* knockdown by shRNA in cancer cell lines from indicated tissue-of-origin including kidney (red) from the DepMap database. Tissue abbreviations are the same as in panel C. **F.** Confocal images of 786-O, 769-P, and BFTC909 cells treated with ML210 and indicated concentrations of vehicle (DMSO) or Lip-1 for the indicated time periods. Nine fields per treatment condition were aligned together for effective visualization. Scale bars in the images represent 50μm. **G.** Immunoblot analysis of HIF-2α protein levels in CCLF_KIPA_0001_N (normal), CCLF_KIPA_0001_T (tumor), and CCLF_KIPA_0002_T (tumor) cells. β-Actin was used as a loading control. Representative plot of experiments repeated three times.

**Figure S2.**
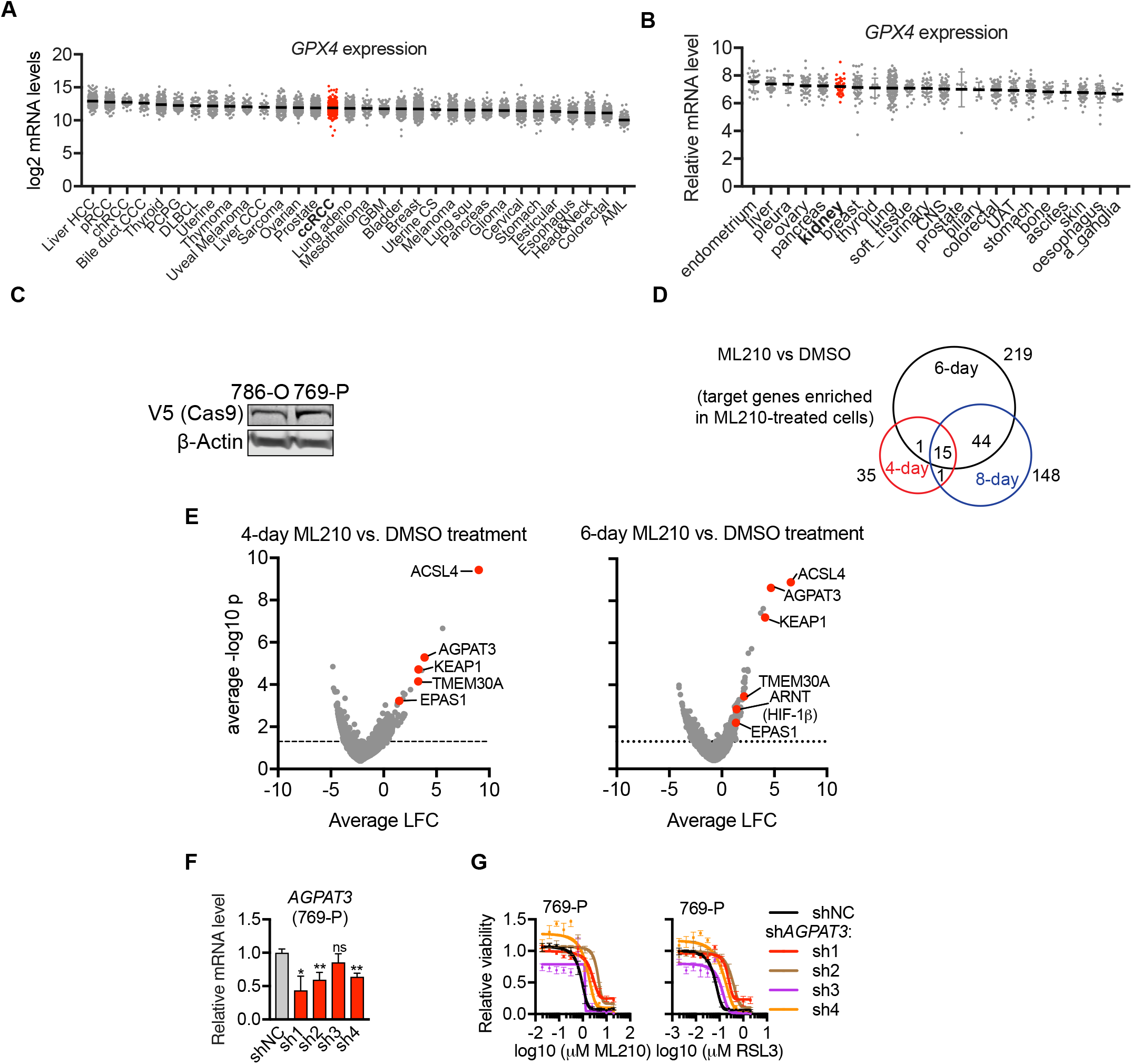
Systematic identification of mediators for ferroptosis susceptibility in ccRCC cells through a genome-wide CRISPR resistance screen. Related to Figure 2. **A.** Expression of *GPX4* mRNA levels in the indicated cancer types including ccRCC (red) from The Cancer Genome Atlas (TCGA) database. Processed data were retrieved from cBioPortal. Log2 RPKM is presented for each sample. The cancer types are ordered by the mean of GPX4 expression. Cancer type abbreviations: Breast, Breast Invasive Carcinoma; DLBCL, Lymphoid Neoplasm Diffuse Large B-cell Lymphoma; Head&Neck, Head and Neck Squamous Cell Carcinoma; Liver HCC, Liver Hepatocellular Carcinoma; Liver CCC, Cholangiocarcinoma; Ovarian, Ovarian Serous Cystadenocarcinoma; Uterine CS, Uterine Carcinosarcoma; Glioma, Brain Lower Grade Glioma; Colorectal, Colorectal Adenocarcinoma; ccRCC, Kidney Renal Clear Cell Carcinoma; Lung adeno, Lung Adenocarcinoma; Prostate, Prostate Adenocarcinoma; Cervical, Cervical Squamous Cell Carcinoma and Endocervical Adenocarcinoma; Bladder, Bladder Urothelial Carcinoma; Esophagus, Esophageal Carcinoma; Pancreas, Pancreatic Adenocarcinoma; PCPG, Pheochromocytoma and Paraganglioma; GBM, Glioblastoma Multiforme; Thyroid, Thyroid Carcinoma; Uterine, Uterine Corpus Endometrial Carcinoma; Melanoma, Skin Cutaneous Melanoma; Stomach, Stomach Adenocarcinoma; Testicular, Testicular Germ Cell Cancer; pRCC, Kidney Renal Papillary Cell Carcinoma; AML, Acute Myeloid Leukemia; chRCC, Kidney Chromophobe; Lung squ, Lung Squamous Cell Carcinoma. **B.** Expression of relative *GPX4* mRNA levels in cancer cell lines from each indicated tissue of origin including kidney (red) characterized in CTRP. The cancer types are ordered by the mean of GPX4 expression. Abbreviations for tissue of origin: CNS, central nervous system; UAT, upper aerodigestive tract; a_ganglia, autonomic ganglia. **C.** Immunoblot analysis confirming the expression of Cas9-V5 protein in 786-O and 769-P cells transduced with constitutive Cas9 expression. β-Actin was used as a loading control. **D.** Venn Diagram summarizing the number of genes that are significantly enriched under each combination of screening conditions. **E.** Volcano plot highlighting (red) the top enriched CRISPR hits comparing 4-day or 6-day ML210 treated 786-O cells with the DMSO-treated condition. LFC, log2 fold change (ML210/DMSO). **F.** qRT-PCR analysis of relative *AGPAT3* mRNA expression in 769-P cells expressing shNC or AGPAT3-targeting shRNAs. *, p<0.05; **, p<0.01. *B2M* was used as a loading control. **G.** Viability curves for shNC or AGPAT3-targeting shRNA-expressing 769-P cells under indicated concentrations of ML210 or RSL3. Viability under each condition is relative to that of the respective DMSO-treated condition. Representative plot of experiments repeated three times. Each data point has four biological replicates, and error bars represent ±S.D.

**Figure S3.**
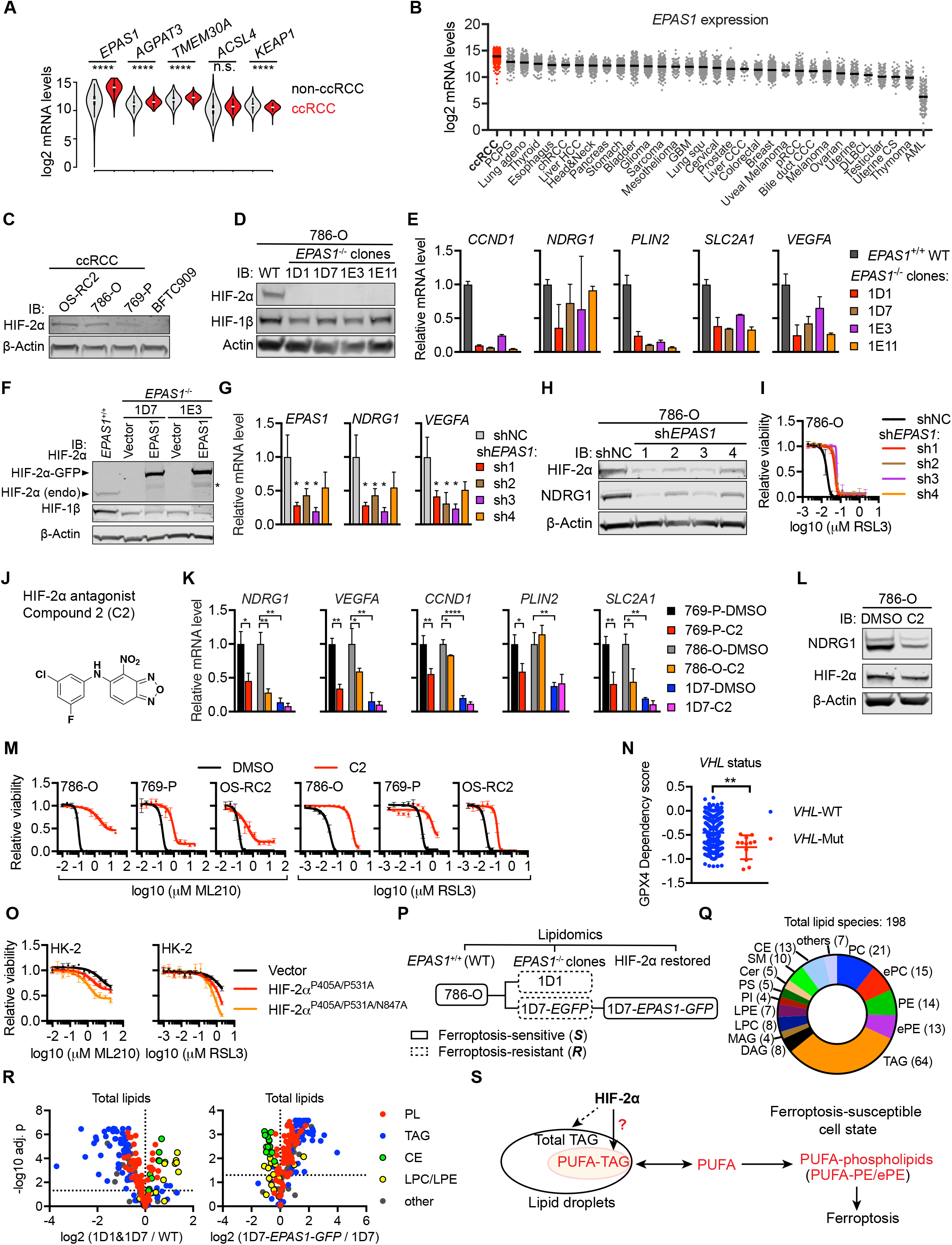
HIF-2α activation is required for GPX4 dependency and ferroptosis susceptibility in ccRCC. Related to Figure 3. **A.** mRNA expression of the indicated genes in ccRCC (N=535; red) or non-ccRCC (N=9,187; grey) tumor samples collected in the TCGA RNA-Seq database. Student’s T-test, ****, p<0.0001; ns, not significant. **B.** *EPAS1* mRNA expression in the indicated cancer types including ccRCC (red) from the TCGA RNA-Seq datasets. Abbreviations are the same as in Figure S2A. **C.** Immunoblot analysis of HIF-2α expression in ccRCC cell lines OS-RC2, 786-O, 769-P, and RTCC cell line BFTC909. β-Actin was used as a loading control. **D.** Immunoblot showing HIF-2α and HIF-1β protein levels in wildtype (WT) 786-O cells and four *EPAS1^−/−^* clones generated by CRISPR/Cas9 and single-cell clone isolation. **E.** qRT-PCR analyses of mRNA levels of HIF-2α target genes *CCND1, NDRG1, PLIN2, SLC2A1* (GLUT1), and *VEGFA* in 786-O WT or the indicated *EPAS1^−/−^* clones. *B2M* was used as an internal control. Error bars represent ±S.D. **F.** Immunoblot analysis of HIF-2α and HIF-1β protein levels in wildtype *(EPAS1^+/+^),* two *EPAS1^−/−^* single-cell clones 1D7 and 1E3, each expressing an empty vector or *EPAS1*-GFP *(EPAS1)* construct. **G.** qRT-PCR analyses of mRNA levels of *EPAS1* and HIF-2α target genes *NDRG1* and *VEGFA* in 786-O cells expressing shNC or EPAS1-targeting shRNAs. *GAPDH* was used as an internal control. Error bars represent ±S.D. Student’s T-test, *, p<0.05. **H.** Immunoblot analysis of NDRG1 and HIF-2α protein expression in 786-O cells expressing shNC or EPAS1-targeting shRNAs. β-Actin was used as a loading control. **I.** Viability curves for 786-O cells expressing shNC or EPAS1-targeting shRNAs treated with indicated concentrations of RSL3. Viability under each condition is relative to that of the no RSL3 treatment condition. Representative plot of experiments repeated three times. Each data point has four biological replicates and error bars represent ±S.D. **J.** Chemical structure of HIF-2α antagonist compound 2 (C2). **K.** qRT-PCR analyses of mRNA levels of HIF-2α target genes *NDRG1, VEGFA, CCND1, PLIN2,* and *SLC2A1* (GLUT1) in 769-P, 786-O-WT and *EPAS1^−/−^* 1D7 cells treated with DMSO or C2 for 3 days. Expression is relative to each of the DMSO-treated WT cells. *B2M* was used as an internal control. Student’s T-test, *, p<0.05, **, p<0.01, ****, p<0.0001, ns, not significant. **L.** Immunoblot analysis of NDRG1 and HIF-2α protein expression in 786-O cells treated with DMSO or C2. β-Actin was used as a loading control. **M.** Viability curves for the indicated cell lines pre-treated with DMSO or C2 for 3 days, then treated together with indicated concentrations of ML210 or RSL3. Viability under each condition is relative to that of the no ML210 or RSL3 treatment condition. Representative plot of experiments repeated three times. Each data point has four biological replicates, and error bars represent ±S.D. **N.** Scatterplot showing the GPX4 dependency scores (CERES) by CRISPR in cancer cell lines possessing wildtype (WT, N=421; blue) or mutant *VHL* (N=12; red) in the DepMap database. Mann-Whitney-Wilcoxon test, **, p < 0.01. **O.** Viability curves for HK-2 cells expressing empty vector or exogenous HIF-2α^P405A/P531A^, or HIF-2α^P405A/P531A/N847A^ constructs, treated with indicated concentrations of ML210 or RSL3. Viability is relative to that of the DMSO-treated condition. Each data point has four biological replicates, and error bars represent ±S.D. **P.** Scheme summarizing the lipidomics experiment with 786-O WT, *EPAS1^−/−^* derivative clones 1D1 and 1D7 (1D7-EGFP), and *1D7-EPAS1-GFP* cells. Three biological replicates were included for each condition. **(S)**, ferroptosis-sensitive; **(R)**, ferroptosis-resistant. **Q.** Pie chart summarizing the number of lipid species from each class detected in the lipidomic profiling of 786-O WT cells. Abbreviations: CE, cholesterol ester; Cer, ceramide; MAG, monoacylglycerol; DAG, diacylglycerol; TAG, triacylglycerol; LPC, lysophosphatidylcholine; LPE, lysophosphatidylethanolamine; PC, phosphatidylcholine; PE, phosphatidylethanolamine; ePC, (vinyl ether-linked) PC-plasmalogen; ePE, (vinyl ether-linked) PE-plasmalogen; PI, phosphatidylinositol; PS, phosphatidylserine; SM, sphingomyelin. **R.** Volcano plot highlighting the changes of the indicated lipid classes between the indicated groups. Abbreviations are the same as in panel **Q**. **S.** Scheme summarizing the role of HIF-2α in lipid metabolism and ferroptosis susceptibility.

**Figure S4.**
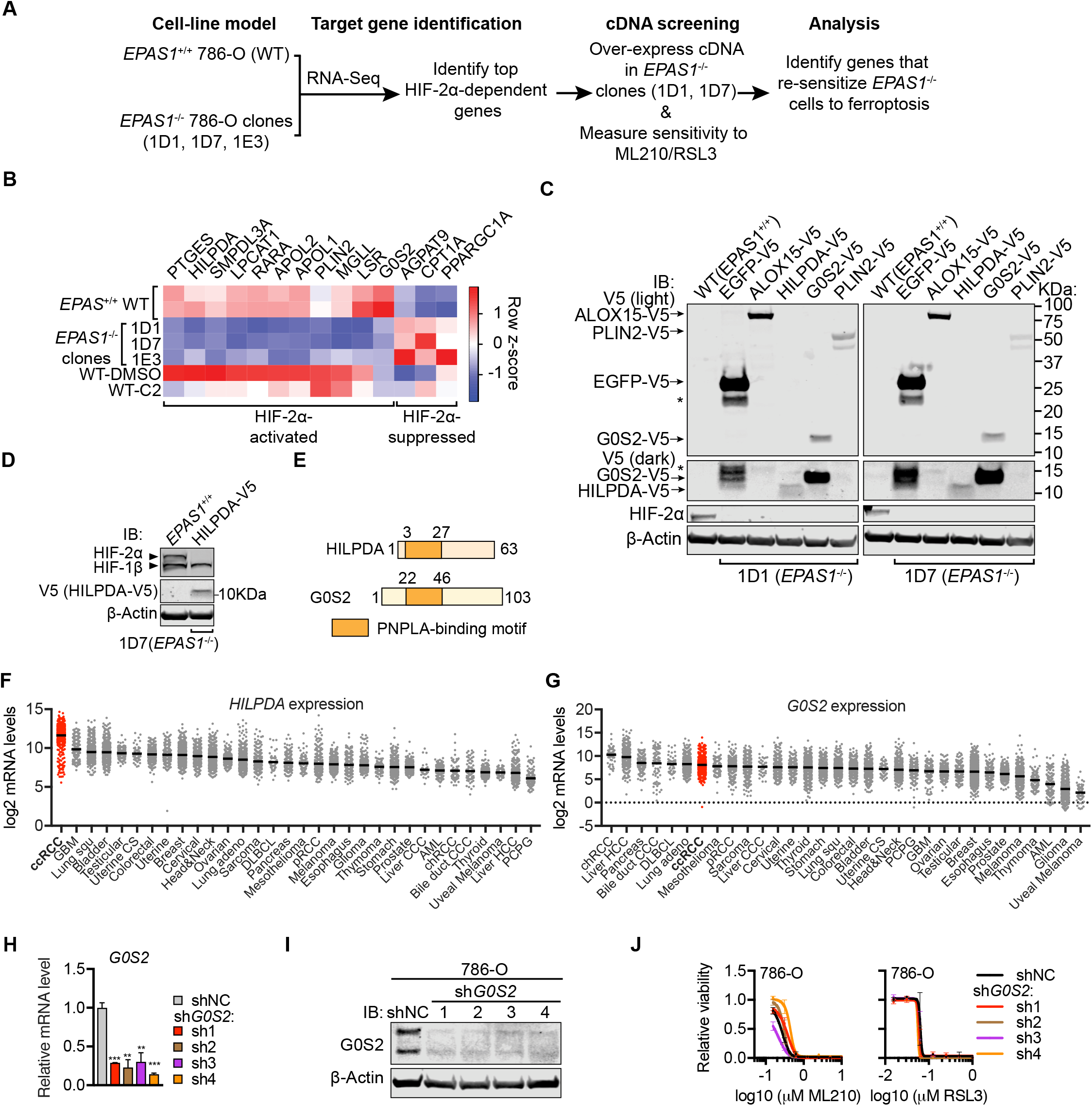
HILPDA and G0S2 mediate HIF-2α’s activity in sensitizing ccRCC cells to GPX4 inhibition-induced ferroptosis. Related to Figure 4. **A.** Scheme summarizing the experimental strategy for identifying the HIF-2α target genes mediating ferroptosis susceptibility in 786-O cells. Briefly, HIF-2α-dependent genes were first identified by RNA-Seq, then cDNA constructs expressing the genes significantly and positively regulated by HIF-2α were transduced into *EPAS1^−/−^* clones to test their activities in re-sensitizing HIF-2α-depleted cells to ferroptosis-inducing agents. **B.** Heatmap showing RNA-Seq results for the HIF-2α-dependent lipid metabolism genes in 786-O cells. **C.** Immunoblot analysis of protein levels of HIF-2α, and V5 tagged EGFP, ALOX15, HILPDA, G0S2, and PLIN2 in *EPAS^−/−^* 1D1 (left) and 1D7 (right) cells. Two different exposure levels were presented for the lower molecular weight part of the membranes. β-Actin was used as a loading control. Representative plot of experiments repeated three times. Error bars represent ±S.D. **D.** Immunoblot analysis of protein levels of HIF-2α, HIF-1 β, and V5 tagged HILPDA in *EPAS^−/−^* 1 D7 cells to verify the results in panel C. β-Actin was used as a loading control. **E.** Scheme summarizing the secondary protein structure of HILPDA and G0S2, showing the shared PNPLA-inhibitory motif (orange). Numbers indicate amino acid residues. **F.** *HILPDA* mRNA expression in the indicated cancer types including ccRCC (red) from the TCGA RNA-Seq datasets. Abbreviations are the same as in **Figure S2A**. **G.** *G0S2* mRNA expression in the indicated cancer types including ccRCC (red) from the TCGA RNA-Seq datasets. **H.** qRT-PCR analysis of *G0S2* mRNA levels in 786-O cells expressing shNC or *shG0S2s. B2M* was used as an internal control. Each condition includes three biological replicates. Error bars represent ±S.D. Student’s T-test. *, p< 0.05. **I.** Immunoblot analysis of G0S2 protein levels in 786-O cells expressing shNC or *shG0S2s.* β-Actin was used as a loading control. **J.** Viability curves of control or *G0S2* knockdown 786-O cells under indicated concentrations of ML210 or RSL3 for 48h. Viability is relative to the respective DMSO-treated conditions. Each data point has four biological replicates, and error bars represent ±S.D.

**Figure S5.**
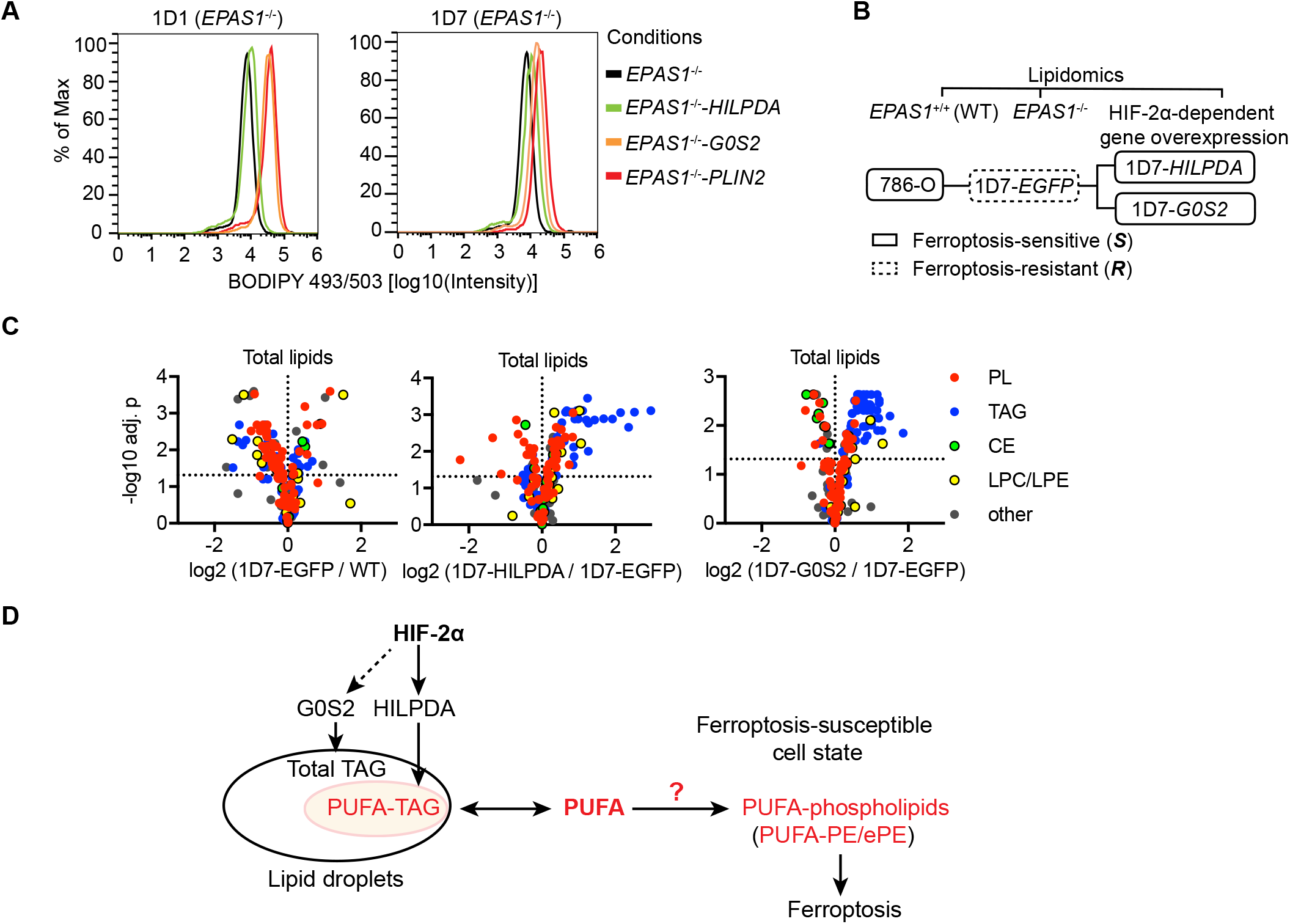
The HIF-2α-HILPDA axis drives the polyunsaturated fatty acyl-lipid-enriched state in ccRCC cells. Related to Figure 5. **A.** Lipid droplet abundances analyzed by flow cytometry quantitation of BODIPY-493/503 signal in 1D1 and 1D7 *EPAS1^−/−^* cells expressing exogenous *EGFP, HILPDA, G0S2,* or *PLIN2.* Representative plot of experiments repeated three times. **B.** Scheme summarizing lipidomic profiling experiments for 786-O and *EPAS1^−/−^* 1D7 cells expressing *EGFP, HILPDA,* or *G0S2.* Three biological replicates were included for each condition. **(S)**, ferroptosis-sensitive; *(**R**),* ferroptosis-resistant. **C.** Volcano plots highlighting the changes of the indicated lipid classes between the indicated groups. Lipid class abbreviations are the same as in **Figure 3H**. **D.** Summary scheme for the HIF-2α-dependent molecular network that drives GPX4 dependency and ferroptosis susceptibility in ccRCC tumors.

**Figure S6.**
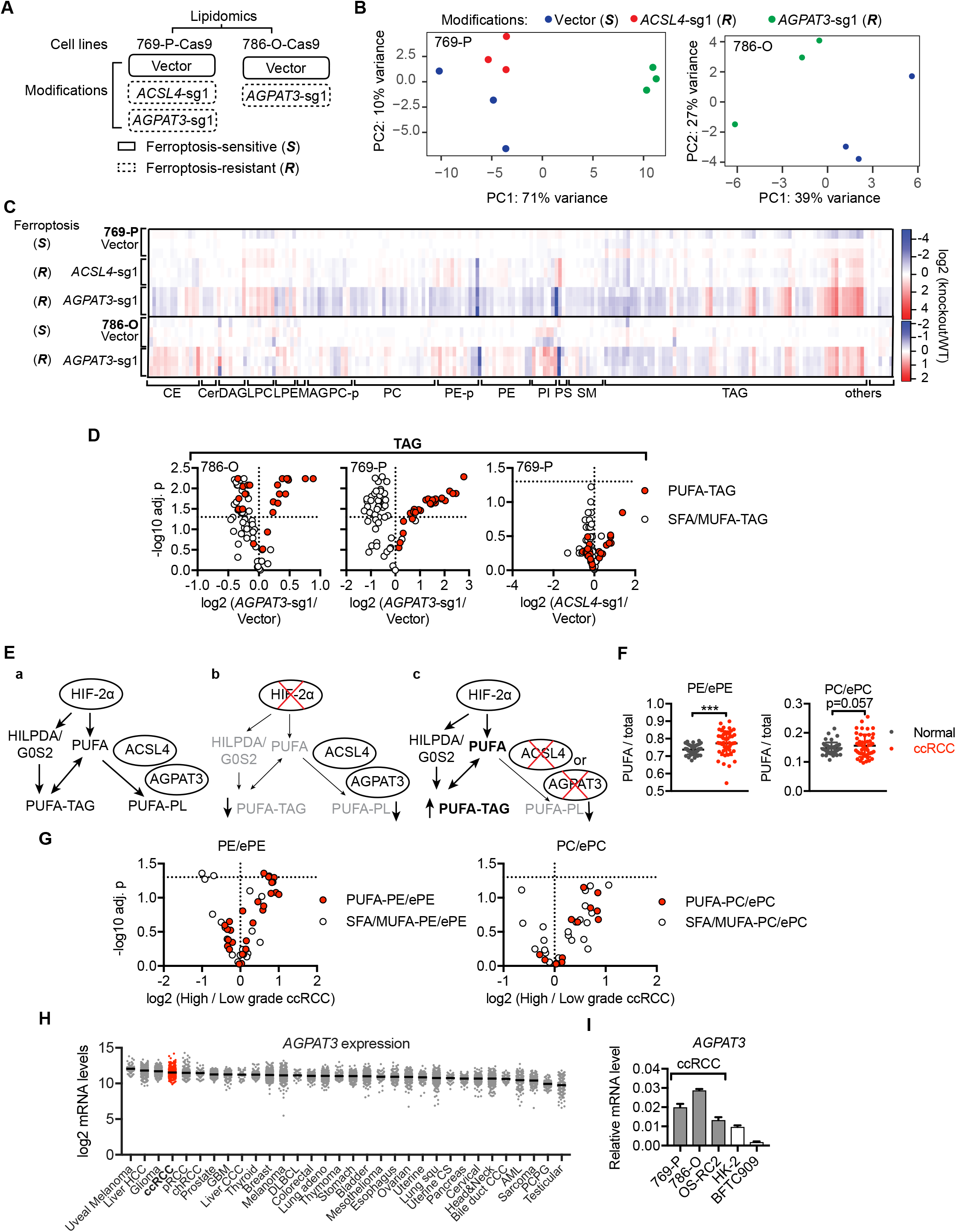
HNF-1β-driven AGPAT3 selectively synthesizes polyunsaturated fatty acyl-phospholipids and primes the renal lineage for GPX4 dependency. Related to Figure 6. **A.** Scheme summarizing lipidomic profiling experiments with 769-P-Cas9-vector, *ACSL4-sg1*, or *AGPAT3-sg1* cells and 786-O-Cas9-vector, *AGPAT3-sg1* cells. Each condition includes three biological replicates. **(S)**, ferroptosis-sensitive; **(R)**, ferroptosis-resistant. **B.** Principal component plots for the lipidomic profiles of the 769-P-Cas9-vector, *ACSL4-sg1* and *AGPAT3-sg1* cells (left), and for the 786-O-Cas9-vector, *AGPAT3-sg1* cells (right). **C.** Heatmap presenting relative lipid abundances in the indicated cell lines. The lipid ratios between ACSL4-sg1, *AGPAT3-sg1* and WT of 769-P and 786-O cells were log2 transformed and plotted. Each group includes three biological replicates. Color key: blue: down-regulated in the test cell line, red: up-regulated in the test cell line. Lipid class abbreviations are the same as in **Figure 3H**. **D.** Volcano plots showing changes in each TAG species grouped as PUFA-TAGs and SFA/MUFA-TAGs between the indicated cell lines. **E.** Scheme summarizing the flow of fatty acid metabolism between free PUFA, PUFA-TAG, and PUFA-phospholipids (PUFA-PL) (a), and major lipidomic changes in HIF-2α-depleted (b), or ACSL4 or AGPAT3-depleted (c) cells. Loss of HIF-2α leads to down-regulation of HILPDA/G0S2 and free PUFA levels, which subsequently reduces PUFA-TAG and PUFA-PL deposition (b). When ACSL4 or AGPAT3 is depleted, PUFA-phospholipid synthesis is blocked and free PUFAs are incorporated into the reciprocal PUFA-TAGs (c). **F.** The ratios between PUFA-PE/ePE and total PE/ePE (left), and between PUFA-PC/ePC and total PC/ePCs in ccRCC tumor samples (N=49; red) and the matched normal tissues (N=49; grey) from previously reported lipidomics datasets. Student’s T-test, ***, p < 0.001. **G.** Volcano plots showing changes in each PE/ePE species grouped as PUFA-PE/ePEs (red fill) and SFA/MUFA-PE/ePEs (white fill) between high-grade (grade III/IV, N=18) and low-grade (grade I/II, N=16) ccRCC tumor samples. **H.** *AGPAT3* mRNA expression in the indicated cancer types including ccRCC (red) from the TCGA RNA-Seq datasets. Abbreviations are the same as in **Figure S2A**. **I.** qRT-PCR analysis of *AGPAT3* mRNA levels in ccRCC cell lines 769-P, 786-O, OS-RC2, immortalized normal renal epithelial cell line HK-2 and RTCC cell line BFTC909. Expression relative to *B2M* was plotted. Representative plot of experiments repeated three times. Each condition includes three biological replicates. Error bars represent ±S.D.

## MATERIALS AND METHODS

### Cell culture

786-O, 769-P, and OS-RC2 cells were cultured in RPMI-1640 (Gibco) media; RCC10RGB, BFTC909, and HEK-293T cells were cultured in DMEM (Gibco) media. Both RPMI-1640 and DMEM media were supplemented with 10% fetal bovine serum and 1% penicillin/streptomycin. HK-2 cells were cultured in keratinocyte serum-free medium (Gibco) supplemented with 0.05mg/mL bovine pituitary extract and 5ng/mL human recombinant epidermal growth factor. All cells were cultured in a humidified incubator at 37°C with 5% CO2. All cancer cell lines were obtained from the Cancer Cell Line Encyclopedia distributed by the Broad Institute Biological Samples Platform and HK-2 cells were obtained from American Type Culture Collection (ATCC). All cells were regularly tested for mycoplasma contamination and cells used in experiments were negative for mycoplasma.

### Compound sources, synthesis, and treatment

ML210 and RSL3 were synthesized according to previously described protocols [28, 42]. The concentration range for ML210 was 0.01953-20μM in 11-concentration experiments and 0.07813-5μM in 7-concentration viability curve measurements. The concentration range for RSL3 is 0.001953-2μM in 11-concentration experiments and 0.01563-1 μM in 7-concentration viability curve measurements. Liproxstatin-1 (Lip-1, Sigma-Aldrich SML1414) was used at 0.5-1 μM. Ferrostatin-1 (Fer-1, Sigma-Aldrich, SML0583) was used at 1-5μM. HIF-2α antagonist compound 2 (C2, Sigma-Aldrich, SML0883) was used at 5-10μM.

### Gene-expression analysis

Total RNA was extracted from cells using RNeasy Mini kit (Qiagen) following the manufacturer’s instructions. cDNA was synthesized using the ProtoScript First Strand cDNA Synthesis kit (New England Biolabs). Quantitative PCR reaction mixtures were prepared with the SYBR Green PCR Master Mix (Thermo Fisher Scientific, Applied Biosystems). The PCR reactions were performed and analyzed on a LightCycler 480 Instrument (Roche). Each sample condition contains at least three biological replicates and all measurements were performed with four technical replicates. Mean and S.D. of biological replicates were presented. Gene-expression levels were normalized to internal control genes including *B2M* and *GAPDH* and was relative to non-treatment condition unless otherwise indicated.

### Immunoblotting

Cells were briefly washed twice with PBS and lysed with 1% SDS lysis buffer containing 10mM EDTA and 50mM Tris-HCl, pH8.0. Lysates were collected, incubated at 95°C for 10min and the protein concentrations were determined by BCA Protein Assay kit (Pierce). Calibrated samples were diluted with 4x LDS sampling buffer (Novus), separated by SDS-PAGE using NuPAGE 4-12% Bis-Tris protein gels (Novus), and transferred to nitrocellulose or PVDF membranes by iBlot2 protein-transfer system (Thermo Fisher Scientific). Membranes were blocked with 50% Odyssey blocking buffer (LiCor) diluted with 0.1% Tween-20-containing TBS (TBST) and immunoblotted with antibodies against TMEM30A (Abcam, ab105062), GPX4 (Abcam, ab41787), ACSL4 (Abcam, ab155282), NDRG1 (Abcam, ab124689), KEAP1 (D6B12, Cell Signaling Technologies, #8047), NRF2 (D1Z9C, Cell Signaling Technologies, #12721), V5 tag (D3H8Q, Cell Signaling Technologies, #13202), HIF-1a (D5F3M, Cell Signaling Technologies, #79233), HIF-1β/ARNT (D28F3, Cell Signaling Technologies, #5537), HIF-2α (D9E3, Cell Signaling Technologies, #7096), β-Actin (8H10D10, Cell Signaling Technologies, #3700) and β-Actin (13E5, Cell Signaling Technologies, #4970). Membranes were then washed with TBST and incubated with IRDye 800CW goat-anti-Rabbit or 680RD donkey-anti-Mouse secondary antibodies (LiCor). Immunoblotting images were acquired on an Odyssey equipment (LiCor) according to the manufacturer’s instructions. All antibodies were diluted at 1:1000 for immunoblotting. Unless otherwise indicated, β-Actin was used as a loading control.

### CRISPR/Cas-mediated genome editing and RNA interference

For CRISPR/Cas9-mediated genome-editing, 786-O and 769-P cells were engineered for Cas9 expression with the pLX-311-Cas9 vector (Addgene 96924), which contains the blasticidin S-resistance gene driven by the SV40 promoter and the SpCas9 gene driven by the EF1 a promoter. sgRNA sequences were cloned into the pLV709 doxycycline-inducible or pXPR_BRD050 constitutive sgRNA expression vectors. For shRNA-mediated RNA interference, shRNAs targeting the genes of interest were pre-cloned into constitutive shRNA expression vectors pLKO.1 or pLKO-TRC005 by the Broad Institute Genetic Perturbation Platform. Lentiviruses were generated from sgRNA/shRNA constructs in HEK293T packaging cells in 96-well plate format using FUGENE6 (Promega) as the transfection reagent and infected 786-O-Cas9 or 769-P-Cas9 for sgRNAs, and 786-O WT cells for shRNAs. Infected cells were selected with puromycin at 2μg/mL starting 48 hours post-infection and propagated for further analysis. Cells transduced with inducible sgRNA constructs were treated with 1μg/ml doxycycline (Sigma-Aldrich) for 7-14 days prior to gene-knockout validation by immunoblot analysis.

### Single-cell cloning

786-O-Cas9 cells were transfected with ribonucleoprotein (RNP) complex containing EnGen Cas9 NLS, S. *pyogenes* (New England Biolabs), Alt-R CRISPR tracrRNA and Alt-R CRISPR crRNA (Integrated DNA Technologies) according to the manufacturer’s instructions (Integrated DNA Technologies) using Lipofectamine RNAiMAX transfection reagent (Thermo Fisher Scientific). Transfected cells were sorted into 96-well plates at 1 cell/well on SONY SH800 cell sorter (SONY). Cells were allowed to grow for 7 days to become single-cell clones and analyzed by immunoblot for effective gene knockout and further analysis. The names of each clone were designated by the original plate number and well position.

### Genome-wide CRISPR screen and data analysis

Pooled lentiviruses for the sgRNA library was prepared using HEK293T cells in T175 flasks as previously described [109]. Prior to screening, the viral titer and volume was pre-determined with pilot experiments to ensure about 30% infection rate in the screening experiment and caution was taken to minimize multiple constructs transduced into the same cell. Optimized surface area for cell growth was pre-determined in pilot experiments to avoid over confluence during screening. Puromycin concentrations were pre-determined with pilot experiments before the screen. For the screening experiment, 150 million 786-O-Cas9 cells were infected with a lentiviral library containing 77,441 sgRNA targeting 18,000 genes in the human genome to ensure each sgRNA is represented by at least 500 cells on average [52]. For the infection, cells added with the calculated lentivirus volume were supplemented with 4μg/mL of polybrene and centrifuged at 2000rpm for 2 hours. Fresh media was added at a 1:1 ratio post-centrifugation. Infected cells were selected with 2μg/mL of puromycin for 96 hours, expanded for another 4 days to reach the desired cell number and split for DMSO or 5μM ML210 treatment. At least 40 million cells were used for each treatment condition to keep the minimum presentation number for each sgRNA above 500. Cells were exposed to ML210 treatment for 4, 6, or 8 days before being cultured in drug-free media for recovery and expansion for 24h. Genomic DNA from cell pellets was purified using the QIAamp DNA Blood Maxi/Midi/Mini kits (Qiagen) according to the manufacturer’s protocols and quantified using a Nanodrop 2000 (Thermo Fisher Scientific).

The sequencing library was prepared, sequenced, and analyzed as previously described [109, 110]. Briefly, sgRNA cassettes were PCR-amplified and barcoded with sequencing adaptors utilizing ExTaq DNA Polymerase (Clontech). PCR products were purified with Agencourt AMPure XP SPRI beads (Beckman Coulter A63880) according to the manufacturer’s instructions, quantified using a Nanodrop 2000, pooled into a master sequencing pool, and sequenced on a MiSeq sequencer (Illumina) with 300nt single-end reads, loaded at 60% with a 5% spike-in of PhiX DNA. For CRISPR screen data analysis, the sgRNA sequences were mapped to a reference file containing all SpCas9 sgRNAs in the library and the sgRNA-associated barcodes were counted and mapped to the barcode reference file. The read count matrix was normalized to reads per million reads (RPM) within each condition. Normalized read counts for each sgRNA in the ML210-treated conditions were compared with those in the DMSO-treated condition. Genes represented by 3-10 sgRNAs were included in the differential enrichment analysis. Genes of interest in each ML210-treated condition were required to have at least 2 sgRNAs exhibiting at least 2-fold enrichment with p values < 0.05. Genes that scored in all three ML210-treated conditions were further validated with shRNA-mediated knockdown.

### Imaging analysis

Cells were plated at 5,000 cells per well in a 96-well, Cell Carrier Ultra Microplate (PerkinElmer) in the appropriate cell culture media, supplemented with 1 μM of liproxstatin-1 where indicated and left overnight. Cells were incubated with 0.1% DMSO (same volume of DMSO as samples with a compound), 10 μM ML210, or 10 μM ML210 + 1 μM of liproxstatin-1 for the indicated times. During the last hour of incubation, the media also contained 60 nM DRAQ7 (Abcam), 1μg/mL Hoechst 33342 (Invitrogen-Thermo Fisher Scientific), and 1 μM BODIPY-581/591 C11 (Thermo Fisher Scientific) for live-cell imaging. Cells were imaged at 63x magnification using an Opera Phenix High-Content Screening System (PerkinElmer, Waltham, MA) equipped with 405 nm, 488 nm, 560 nm, and 647 nm lasers. Image analysis was conducted with Harmony software (PerkinElmer). All images (14 images per well) were collected with the same instrument parameters and processed with the same settings to maximize the ability to compare results between conditions.

### CellTiter-Glo assay for viability analysis

For cellular viability assays, cells were seeded in 384-well opaque white tissue culture and assay plates (Corning) at 1000 cells/well. 18-24 hours after seeding, cells were treated with compounds at indicated concentrations for 48-72h. Cellular ATP levels were quantified using CellTiter-Glo (Promega) on a multi-plate reader (Envision). Relative viability was normalized to the respective DMSO-treated condition unless otherwise indicated. For data presentation, the mean and standard deviation for the four biological replicates of each data point in a representative experiment is presented. Sigmoidal non-linear regression models were used to compute the regression fit curves. For plate-based screening, area-under-curve (AUC) value for each regression curve is calculated and normalized to 1 as the total AUC for the concentration ranges of ML210 or RSL3.

### Lipid droplet abundance analysis by flow cytometry

786-O, 769-P cells, and their derivatives were grown at optimal confluence and stained with BODIPY-493/503 (ThermoFisher Scientific, Molecular Probes) at 2μM final concentration for 30min according to the manufacturer’s instructions. Cells were then briefly washed with PBS, trypsinized, collected, stained with Hoechst 33258 for 5 minutes and filtered through a 70μm nylon filter. The resulted post-staining single-cell suspensions were analyzed on a SONY SH800 cell sorter (SONY) according to the manufacturer’s protocols. The filter used for detecting the BODIPY-493/503 signal was FITC 488nm. A minimum of 10,000 cells were analyzed for each condition, and each experiment was independently performed at least twice. Representative experimental results are shown.

### cDNA screening

cDNAs for the HIF-2α-dependent genes identified by RNA-Seq were obtained from the previously described human cDNA library collection at the Broad Institute Genetic Perturbation Platform [111]. These cDNAs were constructed for mammalian expression in the pLX-TRC317 vector system. Lentiviruses were produced with the cDNA constructs in HEK293-T cells in 96-well format. *EPAS1^−/−^* 786-O clones were infected with the cDNA lentivirus array, selected with 2μg/mL puromycin for 96 hours and analyzed for the cells’ ferroptosis susceptibility 7 days post-infection. Control vectors expressing EGFP were used to evaluate the infection rate. Genes of interest were identified as having significantly shifted ML210 and RSL3 sensitivity curves compared with EGFP-expressing *EPAS1^−/−^* 786-O cells. Protein expression of the top hit genes and controls was verified by immunoblot analysis.

### RNA-Seq and data analysis

RNA-Seq analysis was performed with wildtype and three *EPAS1^−/−^* 786-O single-cell clones generated by CRISPR/Cas9 to identify the HIF-2α-responsive genes. Samples from wildtype 786-O cells treated with DMSO or HIF-2α antagonist compound C2 for 3 days [64] were also included in this experiment. Total RNA was extracted from adherent 786-O cells and derivatives using the RNeasy Mini Kit (Qiagen Inc.) according to the manufacturer’s instructions. The RNA sequencing library was prepared using NEB-Next Ultra RNA Library Prep Kit following the manufacturer’s recommendations. Briefly, mRNAs were first enriched with Oligo-d(T) beads and fragmented for 15 minutes at 94 °C. First strand and second strand cDNA were subsequently synthesized, end repaired, and adenylated at 3’ ends. Universal adapter was ligated to cDNA fragments, followed by index addition and library enrichment with limited-cycle PCR. Sequencing libraries were validated using the Agilent Tapestation 4200, and quantified using a Qubit 2.0 Fluorimeter as well as by quantitative PCR. The sequencing libraries were multiplexed and clustered on one lane of a flowcell. After clustering, the flowcell was loaded on the Illumina HiSeq instrument according to the manufacturer’s instructions. The samples were sequenced using a 2×150 paired-end configuration. Image analysis and base calling were conducted by the HiSeq Control Software. Raw sequence data generated from Illumina HiSeq was converted into fastq files and de-multiplexed using Illumina’s bcl2fastq 2.17 software. One mismatch was allowed for index sequence identification.

For RNA-Seq data analysis, raw paired-end 150bp/150bp sequencing reads were mapped to human genome build hg19 using *Bowtie2* (v2.3.1) with standard settings. On average 66% of read pairs were uniquely mapped to the hg19 genome. Mapped reads were counted to gene features by the *htseq-count* function from *HTSeq* (version 0.9.1) with standard settings. The read count matrix was normalized to library size and analyzed for differentially expressed genes with *DESeq2 (Bioconductor).* Heatmaps were generated using the *heatmap.2* function in *gplots* package (The Comprehensive R Archive Network).

### Lipidomic profiling and data analysis

Analyses of polar and non-polar lipids were conducted using an LC-MS system comprising a Shimadzu Nexera X2 U-HPLC (Shimadzu Corp.; Marlborough, MA) coupled to an Exactive Plus orbitrap mass spectrometer (Thermo Fisher Scientific; Waltham, MA). Lipids were extracted from cells with 0.8mL isopropanol (HPLC Grade; Honeywell). Three replicates were analyzed for each cell line or condition. Cell extracts were centrifuged at 10,000 rcf for 10 minutes to removed residual cellular debris prior to injecting 10 μL onto an ACQUITY BEH C8 column (100 x 2.1 mm, 1.7 μm; Waters, Milford, MA). The column was eluted isocratically with 80% mobile phase A (95:5:0.1 vol/vol/vol 10 mM ammonium acetate/methanol/formic acid) for 1 minute followed by a linear gradient to 80% mobile-phase B (99.9:0.1 vol/vol methanol/formic acid) over 2 minutes, a linear gradient to 100% mobile phase B over 7 minutes, then 3 minutes at 100% mobile-phase B. MS data were acquired using electrospray ionization in the positive-ion mode over 200–1100 *m/z* and at 70,000 resolutions. Other MS settings were: sheath gas 50, in source CID 5 eV, sweep gas 5, spray voltage 3 kV, capillary temperature 300°C, S-lens RF 60, heater temperature 300°C, microscans 1, automatic gain control target 1e6, and maximum ion time 100 ms. Raw data were processed using TraceFinder 3.3 (Thermo Fisher Scientific; Waltham, MA) and Progenesis QI (Nonlinear Dynamics; Newcastle upon Tyne, UK) software for detection and integration of LC-MS peaks. Lipid identities were determined based on comparison to reference standards and reference plasma extracts and were denoted by total number of carbons in the lipid acyl chain(s) and total number of double bonds in the lipid acyl chain(s).

For lipidomics data analysis, median normalization was performed between each sample in the same experiment. Differential-abundance analysis was performed between previously annotated lipid species (about 200 lipids were previously annotated) using Student’s T-test. For fold-change analysis, each dataset was normalized to the mean of the WT cell condition for each lipid species, and the ratio between Test/WT was log2 transformed and presented as heatmaps, bar graphs or volcano plots. P values are adjusted for multiple-test correction using Benjamini-Hochberg correction method and presented as “-*log10 p adj.”.* For principal component plots, the log-ratios between Test/WT for each lipid were further median normalized and computed using the principal component analysis function in DESeq2 *(Bioconductor)* using *RStudio.*

### Public database queries

For CTRP datasets (portals.broadinstitute.org/ctrp/), a data matrix containing the normalized AUC values of each compound in each cell lines (https://ocg.cancer.gov/programs/ctd2/data-portal, ftp://caftpd.nci.nih.gov/pub/OCG-DCC/CTD2/Broad/, and https://github.com/remontoire-pac/ctrp-reference/tree/master/auc) was used for the downstream analyses. Compounds that were profiled in at least 2/3 of the total solid cancer cell line collection, and cell lines that were profiled using at least 50% of the compounds, were included in the statistical analysis. Primary cancer types that contain >5 cell lines profiled for compound sensitivities were presented. Mann-Whitney-Wilcoxon tests were performed between cancer cell lines from each cancer type and other sCCL collections. Statistical significances were adjusted for multiple-test correction using the Benjamini-Hochberg correction method. For gene-expression analysis of TCGA tumor samples, normalized RNA-Seq results for each gene of interest was downloaded through cBioPortal (www.cbioportal.org) and replotted by ranking the mean of log2-transformed expression levels (RPKM) in each cancer type. For Cancer Dependency Map (DepMap) datasets, dependency scores for each gene of interest, including CERES scores from CRISPR/Cas9 screening experiments and ATARis scores for RNA interference screening experiments, were downloaded from the DepMap web portal (https://depmap.org/portal). ChIP-Seq datasets for Hnf-1β in mIMCD cells (GSE71250) and for HIF-2α/HIF-1β in 786-O cells (GSE34871) were downloaded from Gene Expression Omnibus (https://www.ncbi.nlm.nih.gov/geo/).

### Animal staudies

All animal experiments were approved by the Institutional Animal Care and Use Committee (IACUC) of the Broad. Briefly, 3-4-week-old, male athymic nude mice were used for 786-O xenograft experiments. 786-O cells and derivatives were engineered with luciferase and EGFP expression *in vitro* prior to implantation. Five million cells for each injection were trypsinized, resuspended in 50μl PBS containing 0.5μM DMSO or Lip-1, mixed with 50μl Matrigel, and implanted to the subcutaneous space of mice. Tumor volumes were monitored and measured on a weekly basis. For Lip-1 treatment, Lip-1 was first dissolved in DMSO then diluted with PBS and injected into mice at 20 mg/kg body weight daily. Lip-1 treatment was continued for 10 days before withdrawal. For luciferase imaging, each mouse was injected with Xenolight D-luciferin (PerkinElmer) at 150mg/kg and imaged with an IVIS Spectrum Xenogen equipment (Caliper Life Sciences). Tumor volumes were quantified by measuring the length (L) and width (W) of the tumor using a caliper and calculated according to *V=(L*W*W)/2.*

### Patient-derived cancer cell model generation

Human kidney renal cell carcinoma samples, CCLF_KIPA_0001_T and CCLF_KIPA_0002_T, were obtained from patients, with their informed consent, at the Dana Farber Cancer Institute (DFCI), and all procedures were conducted under an Institutional Review Board (IRB)-approved protocol. Patient tumor resections were placed in a sterile conical tube containing DMEM media (Thermo Fisher Scientific, cat. #11995073) with 10% FBS (Sigma Aldrich, cat. #F8317), 1% penicillin-streptomycin (Thermo Fisher Scientific, cat. #15140163), 10μg/ml of gentamicin, and 250 ng/ml fungizone on wet ice during transport from the operating room to the research laboratory. Resections were placed in a 15ml conical flask with 5ml DMEM media, 10% FBS, 1 % penicillin-streptomycin, and the digestion enzymes regular collagenase 1 ml (StemCell, cat. #07912) and dispase 1 ml (StemCell Technologies, cat. #07913). The flask was placed on a rotator and incubated at 37°C for 1 hour. The cells were then centrifuged at 1,000 rpm for 5 min. Cell pellets were resuspended in a 50:50 mix culture medium of Smooth Muscle Growth Medium-2 (Lonza CC-3182) and ACL4 media with 5% FBS [112, 113]. Later, suspended cells were plated into a 96-well plate. The medium was changed every 3 days, and cells were maintained at 37°C in a humidified 5% CO2 incubator. RCC cells were passaged using Gibco TrypLE Express (Thermo Fisher Scientific, cat. # 12604039) to detach cells when the cells reached 80–90% confluence. CCLF_KIPA_0001_N cells were prepared with a similar protocol but cultured with conditional media as previously described [114].

### Statistical analysis

Data are generally expressed as mean±S.D. unless otherwise indicated. No statistical methods were used to predetermine sample sizes. Statistical significance was determined using a two-tailed Mann-Whitney-Wilcoxon test or Student’s T-test using Prism 7 software (GraphPad Software) unless otherwise indicated. The Benjamini-Hochberg correction method was used to adjust the p values where multi-testing corrections were involved. Statistical significance was set at p<0.05 unless otherwise indicated.

